# Conservation of southern yellow-cheeked gibbons (*Nomascus gabriellae*) in the Anthropocene

**DOI:** 10.1101/2025.01.02.631065

**Authors:** Pavla Piskovska, Vit Piskovsky, Susan Cheyne

## Abstract

The endangered southern yellow-cheeked gibbons (*Nomascus gabriellae*) occupy a fragmented habitat in Vietnam and Cambodia, with declining populations due to hunting, habitat loss and climate change. This study integrates population viability modelling with expert surveys to offer comprehensive insights into the species’ current state and its future prospects in a human-shaped world. Our modelling confirms the current population decline and suggests that a reduction in habitat loss and hunting is necessary to save this unique species from extinction. While declining habitat poses a long-term extinction risk, the current hunting rates threaten the species’ survival on the timescale of several decades. Crucially, large populations in Phnom Prich and Keo Seima Wildlife Sanctuaries and Cat Tien, Chu Yang Sin and Bu Gia Map National Parks play a key role in the survival of this species, as does the connectivity of these habitats. The expert surveys confirm the priority of protecting large populations and provide specific conservation actions for the case of Cat Tien National Park that are based on systematic stakeholder mapping. Altogether, this work assesses the population viability of southern yellow-cheeked gibbons in the Anthropocene and suggests conservation strategies that incorporate both ecological and social aspects.

**Graphical Abstract:** 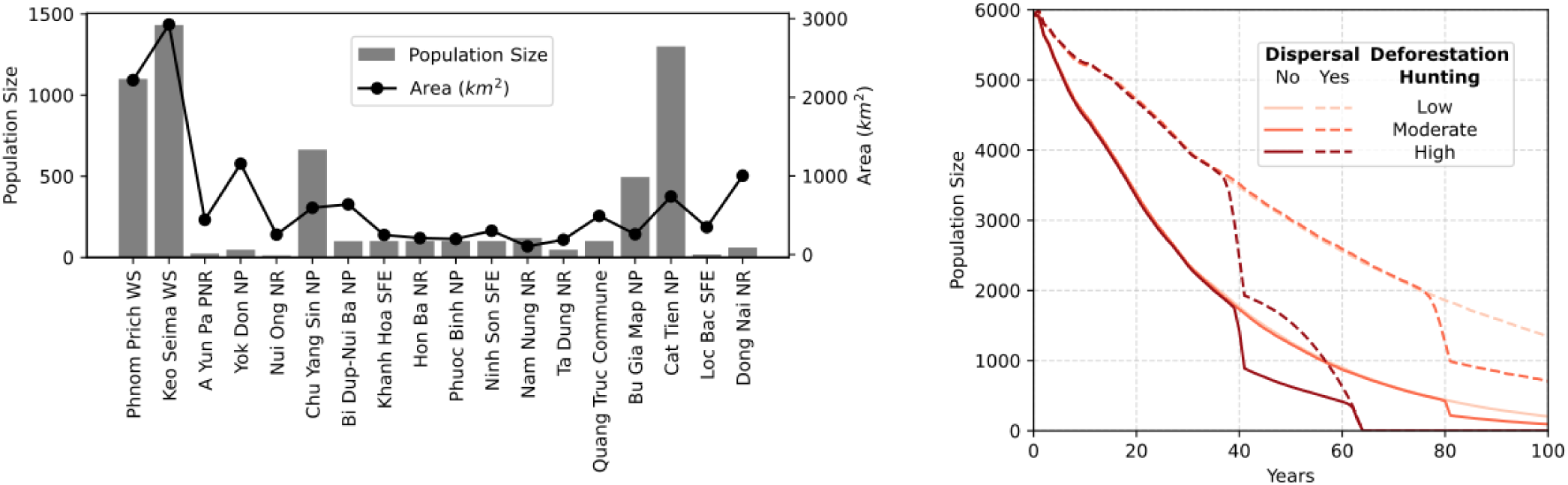

**Highlights:**

1. Southern yellow-cheeked gibbons are at risk of extinction due to hunting, habitat loss and degra- dation.
2. Habitat loss poses a long-term risk while hunting endangers southern yellow-cheeked gibbons on the timescale of the following few decades.
3. Protecting large populations in Phnom Prich and Keo Seima Wildlife Sanctuaries and Cat Tien, Chu Yang Sin and Bu Gia Map National Parks plays a key role in saving southern yellow-cheeked gibbons from extinction.
4. Specific conservation actions based on a systematic stakeholder mapping are identified for Cat Tien National Park.

## 1. Introduction

Resolving the complex relationship between humanity and the natural world is a key challenge in biological conservation. In the current epoch, sometimes referred to as the Anthropocene (Lewis and Maslin, 2015), the anthropogenic activities have resulted in widespread habitat loss, pollution, and climate change, threatening the survival of numerous species (Steffen et al., 2018; Mahmoud and Gan, 2018; Scanes, 2018; Prakash and Verma, 2022; Harfoot et al., 2021) and creating a challenging environment for their conservation (Collard et al., 2015; Malhi et al., 2014). For instance, habitat loss and fragmentation caused by deforestation, agriculture and urbanization have led to a decline in suitable habitats for many species (Hanski, 2011; Scanes, 2018; Haddad et al., 2015). In these fragmented habitats, the native populations are exposed to human-induced stresses, such as imported invasive species (Didham et al., 2007; Hansen et al., 2018; Resasco et al., 2014), pollution (Sigmund et al., 2023; Maiti and Chowdhury, 2013), hunting (Azhar et al., 2013; Benitez-Lopez et al., 2017) and wildlife trade (Morton et al., 2021; Symes et al., 2018), pushing them further towards extinction.

Among the species significantly impacted by these human-induced pressures are primates (Almeida-Rocha et al., 2017). Logging reduces their habitat, which is further fragmented by roads that increase their road kills and allow the poachers to access their habitat and hunt them as food, pests or for illegal pet trade (Lappan and Ruppert, 2019; Gates, 1996; Brodie et al., 2015; Rimbach et al., 2013). Our closest-related primates, the apes, are particularly affected by the Anthropocene (Caldecott and Miles, 2005; Cheyne et al., 2023). For example, their genetic similarity makes them susceptible to human pathogens that are easily transmitted in their fragmented habitats (Dunay et al., 2018). As a result, both great apes (chimpanzees, gorillas, and orangutans) and lesser apes (also known as gibbons) are highly endangered (IUCN, 2023) and require dedicated conservation efforts (Junker et al., 2021; Cowlishaw and Dunbar, 2021). The conservation of gibbons is particularly important (Fan and Bartlett, 2017), as gibbons harbour a large diversity of 20 different subspecies with 70% endangered and 25% critically endangered (IUCN, 2023) and some of these subspecies provide key ecosystem services, such as seed dispersal provided by southern yellow-cheeked gibbons (Hai et al., 2018).

This paper will focus on the endangered southern yellow-cheeked gibbons (*Nomascus gabriellae*). Southern yellow-cheeked gibbons inhabit highly fragmented regions in Vietnam and Cambodia (Traeholt et al., 2005; Rawson et al., 2011), facing the dangers of deforestation (Nuttall et al., 2022), illegal hunting (Ibbett et al., 2021) and pet trade (Duy et al., 2010). Correspondingly, their study has commonly focused on different fragmented sites, such as those in Phnom Phrich Wildlife Sanctuary (Channa and Gray, 2009; Gray et al., 2010), Keo Seima Wildlife Sanctuary (Rawson et al., 2009; Pollard et al., 2007), Cat Tien National Park (Kenyon, 2008; Kenyon et al., 2011), Chu Yang Sin National Park (Vu et al., 2016), Ta Dung Nature Reserve (Duc et al., 2010), or Dong Nai Nature Reserve (Dang and Vinh, 2011). Moreover, the studies that integrated information from multiple sites focused separately on the Cambodian (Traeholt et al., 2005) or Vietnamese populations (Traeholt et al., 2005), even though such populations could be connected by dispersal. As a result, there is a need to integrate these studies to guide the global conservation strategies of this unique species. Moreover, further integration of biological studies with the social aspects of nature conservation can be beneficial, as emphasised by the ethno-primatological approach (Fuentes, 2010, 2012; Dore et al., 2017). In particular, deforestation and hunting are manifestations of a complicated human-wildlife conflict that involves multiple stakeholders (Nyhus, 2016; Treves et al., 2006). Moreover, as the southern yellow-cheeked gibbons inhabit two different political countries, there is a need for international cooperation and transboundary conservation (Mason et al., 2020).

In this paper, we will use an integrated ethno-primatological approach to assess the population viability of southern yellow-cheeked gibbons (*Nomascus gabriellae*) and propose global and local conservation strategies that take into account the social contexts of nature conservation. Specifically, we will use population viability modelling (PVM) to predict the extinction risks associated with deforestation and hunting and integrate these predictions with the local knowledge of field experts. PVM is a stochastic mathematical model for estimating extinction risks over a fixed period (Souĺe, 1985; Stark et al., 2012; Marshall et al., 2008) and has been previously used to guide efficient conservation strategies for various gibbon species (Tunhikorn et al., 1994; Molur et al., 2005; Fan et al., 2013; Bryant, 2014; Smith et al., 2017), including several populations of southern yellow-cheeked gibbons in Vietnam (Trae-holt et al., 2005). Moreover, we will use the expert surveys to confront the PVM with experience from the field and extend our study to social contexts that are equally crucial for biodiversity conservation, such as the mapping of stakeholders (Raum, 2018; Sandroni et al., 2022) and human-wildlife conflicts (Nyhus, 2016; Treves et al., 2006).

## 2. Methods

### 2.1. Study species

Southern yellow-cheeked gibbons live predominantly in long-term monogamy (Traeholt et al., 2005), with occasional cases of polygamy (Barca et al., 2016). Males (Traeholt et al., 2005) and females (Fan et al., 2021) are believed to be capable of first reproduction at the age of around 10 years, while the maximum age of reproduction has been estimated at 28 years for females and 30 years for males (Traeholt et al., 2005). Between these age boundaries, females have at most one brood per year due to a gestation period of 7 months (Hien and Binh, 2024) and give birth to a single offspring (Traeholt et al., 2005; Hien and Binh, 2024). In captivity, the offspring is a male with a chance of 52% (Fan et al., 2021). When the offspring is born, it remains dependent on the mother for about 2.2683 *±* 1.2342 years (Fan et al., 2021) and stays in the family of origin until the age of 6-8 years before it separates to start its own family (Hien and Binh, 2024). For more details, see the summary of the species data in Supplementary Table S1.

### 2.2. Study sites

The southern yellow-cheeked gibbons inhabit multiple fragmented sites (Rainer et al., 2015; Rawson et al., 2011). Table 6 shows the areas and population sizes of these site, where we note that several sites (sites 19-23 in Table 6) have unknown population size, while the population sizes of several other sites (sites 3, 6, 16 in Table 6) must have been estimated from group counts based on the assumption that an average group of southern yellow-cheeked gibbons contains 4 individuals (Duc et al., 2010; Geissmann, 2000). Crucially, the carrying capacity of these sites can be estimated from their area, using the assumption that 1 km^2^ can carry 4 individuals (Traeholt et al., 2005).

The knowledge of carrying capacities for each site is particularly helpful for predicting dispersal between different fragmented habitats. Since the dispersal of gibbons is relatively understudied (Brockelman et al., 1998; Chatterjee, 2006), we considered two extreme dispersal scenarios: (a) no dispersal and (b) ideal free dispersal. In the latter scenario, we assume that gibbons disperse between the ages of 7-10 years with a survival probability of 85% (Traeholt et al., 2005). Moreover, we assume that the dispersal is only permitted between neighbouring sites in proportion to their carrying capacities. This assumption is based on the theory of the ideal free distribution, which postulates that individuals disperse in proportion to resources at each site (Tregenza, 1995) and predicts that such dispersal is evolutionarily stable (Krivan et al., 2008).

### 2.3. Threats

Previous studies have suggested that deforestation and hunting for food or pet trade are the two major threats endangering southern yellow-cheeked gibbons (Traeholt et al., 2005; Rawson et al., 2011, 2020; Channa and Gray, 2009; Ibbett et al., 2021), which we focus on in this work. The annual deforestation rate can be estimated from satellite data (World Resources Institute, 2014), with values in Cambodia at -1.25% and in Vietnam at -0.8%. The quantification of hunting is more complicated since hunting is illegal, which creates a complex environment for surveying this human activity (Nuno and St. John, 2015; Ibbett et al., 2021). For this reason, we assume that hunting is similar to a related gibbon species (*Hylobates moloch*) that inhabits fragmented sites with similarly sized populations (Smith et al., 2017). Since infants are most valuable in the illegal pet trade market, it is beneficial for the hunters to target females that carry an infant (Rawson et al., 2011). Therefore, we estimate the annual hunting rate of a female and an infant at 1.26%, which is the average value of *Hylobates moloch* hunting rates reported for three distinct sites in Smith et al. (2017). To correct for the imprecission of these estimates, this work will consider three different scenarios (Table 6): low severity of threats (deforestation and hunting at 50% of the estimated values), moderate severity of threats (estimated values) and high severity of threats (deforestation and hunting at 200% of the estimated values).

### 2.4. Population viability model

We used the species data (Supplementary Table S1), sites data (Table 6) and threats data (Table 6) to construct a population viability model in VORTEX 10.6 (Lacy and Pollak, 2021) where a specification of dispersal and threat levels defines a scenario of this model (see Supplementary Code). For a given scenario, we sampled 100 different realizations of the population dynamics over the next 100 years to calculate the extinction risks. In particular, we distinguished local extinction events (referring to a population of a single site) and global extinction events (referring to the metapopulation of all sites), where extinction is defined by at most one remaining sex. To quantify these extinction events, we measured the stochastic mean growth rate, the probability of extinction and the mean time to first extinction. The stochastic mean growth rate refers to the mean per capita growth rate across all realizations of the population dynamics. The probability of extinction refers to the proportion of realizations where the population goes extinct. Finally, the mean time to first extinction refers to the mean time of the first extinction across all realizations of the population dynamics, where we note that multiple local extinction events are possible if sites can be recolonized from neighbouring populations. The output data of the models can be found in the Supplementary Data.

### 2.5. Expert surveys

In addition to the population viability model, we conducted surveys about the conservation of southern yellow-cheeked gibbons with local field experts, with the goal to ingrate both biological and social aspects of nature conservation. Indeed, similar mixed-methods approach is useful in preserving biological species in habitats shaped by human actions (Dore et al., 2017), such as the habitat of southern yellow-cheeked gibbons. Adhering to the University of Oxford CUREC ethical guidelines, we recruited field experts by personalized emails and social media posts. The surveys were completed online based on a voluntary consent and the collected data were anonymized before further processing.

The surveys consisted predominantly of open questions and were structured into four sections:

- Sites (section 1): In this part, the participants were asked about various features of different sites where the presence of *Nomascus gabriellae* has been reported. The participants were provided with the list of sites from Table 6 and were recommended to use the same site names, but were also encouraged to mention sites that do not appear on this list. The participants were asked to name the first, second and third most important site for the conservation of *Nomascus gabriellae* in Vietnam and Cambodia and explain the factors that affected their choice. They were further asked to name a site where the population of *Nomascus gabriellae* is in the lowest (resp. highest) risk of local extinction and were asked to provide reasons for this risk assessment. The goal of this section was to obtain a comparison between the modelling and expert surveys.
- Stakeholders (section 2): In this part, the participants were asked about stakeholders that affect *Nomascus gabriellae* in a site they are most familiar with, providing the site name, list of stakeholders, their roles and relationships. The goal of this section was to construct a stakeholder mapping (Raum, 2018; Sandroni et al., 2022) and identify any human-wildlife conflicts (Nyhus, 2016; Treves et al., 2006).
- Threats (section 3): In this part, the participants were asked about threats posed to *Nomascus gabriellae* at the site from section 2. The participants were asked to name the first, second and third biggest threat to *Nomascus gabriellae* and explain their reasoning. The goal of this section was to uncover threats ignored by our modelling and to compare their expected impact with the modelling results.
- Conservation actions (section 4): In this part, the participants were asked about conservation actions for *Nomascus gabriellae* at the site from section 2. The participants were asked to name the first, second and third most effective conservation action for *Nomascus gabriellae* and explain their reasoning. The goal of this section was to identify effective conservation actions and see how they address the key threats from section 3.

To analyze the questions that involved ranking of three most important features (sites, threats, conservation actions), the first most important feature was given a score of 3, the second most important feature was given a score of 2 and the third most important feature was given a score of 1. For each feature, the total score was computed by summing all scores in all answers and the average score was computed by dividing the score by the participant number. In this scoring, we labelled the score of 3 as the Top 1 option, the score of 2 as the Top 2 option and the score of 3 as the Top 3 option. It should be noted that due to a small number of field experts (*N* = 3) that focus on the conservation the souther yellow-cheeked gibbons and that consented to taking part in our surveys, we were unable to perform any statistical tests and the results should be read with caution.

## 3. Results

### 3.1. Populations inhabit fragmented habitat

The yellow-cheeked gibbons are distributed in Vietnam and Cambodia within significantly fragmented habitats (Table 6 and Fig. 1). In Cambodia, the habitat seems less fragmented and contains two large sites, Phnom Prich and Keo Seima Wildlife Sanctuaries. These sites constitute 37.4% of the estimated total habitat area (Fig. 1) and 42.7% of the estimated metapopulation size (Fig. 2a, Supplementary Fig. S1). In contrast, the Vietnamese habitat is more fragmented, with Cat Tien and Chu Yang Sin National Parks being the two most populated sites. These sites carry 33.2% of the estimated metapopulation size (Fig. 2a, Supplementary Fig. S1) and constitute 10.8% of the estimated total habitat area (Fig. 1). Bu Gia Map National Park is the third most populated site in Vietnam, which contributes by an additional 8.4% to the estimated total population size (Fig. 2a, Supplementary Fig. S1) and by 2.1% to the estimated southern yellow-cheeked gibbon habitat (Fig. 1). Altogether, these five sites are estimated to carry 84.3% of the estimated metapopulation size, located in 50.3% of the estimated habitat area. This result suggests the population density in the other sites is either lower or not sufficiently monitored (see also Supplementary Fig. S2).

**Figure 1:**
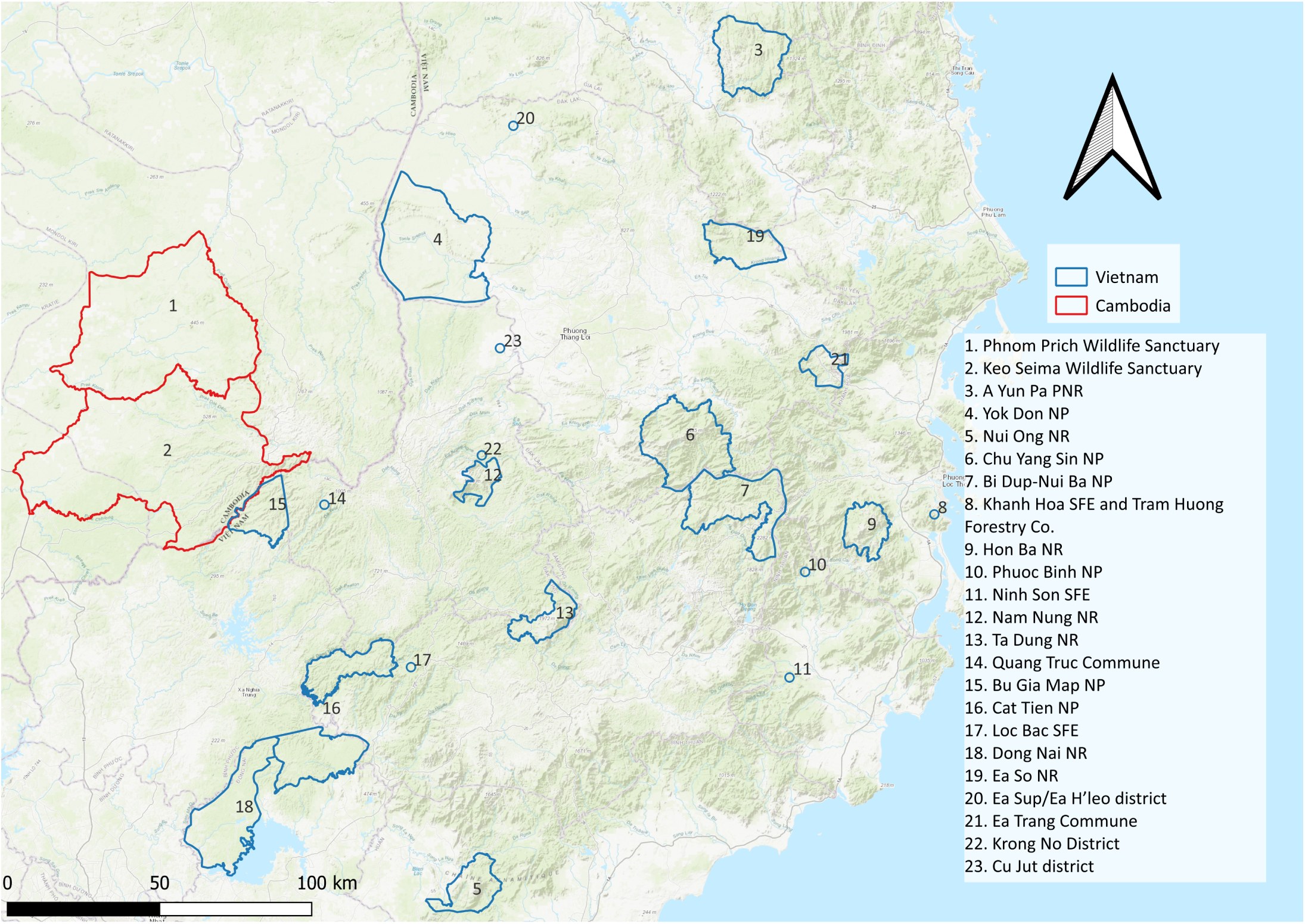
Areas with populations of southern yellow-cheeked gibbons in Vietnam and Cambodia. Areas in Cambodia (1-2, red) and Vietnam (3-23, blue) are represented by polygons (resp. point) for regions with strictly delimited (resp. uncertain) boundaries. The figure was generated in qGIS from The World Topographic Map (ESRI, 2021), National Protected Areas of Vietnam dataset (WCMC, 2021) and Natural protected areas in Cambodia (1993- 2023) dataset (Cambodian Ministry of Environment et al., 2023).

**Figure 2:**
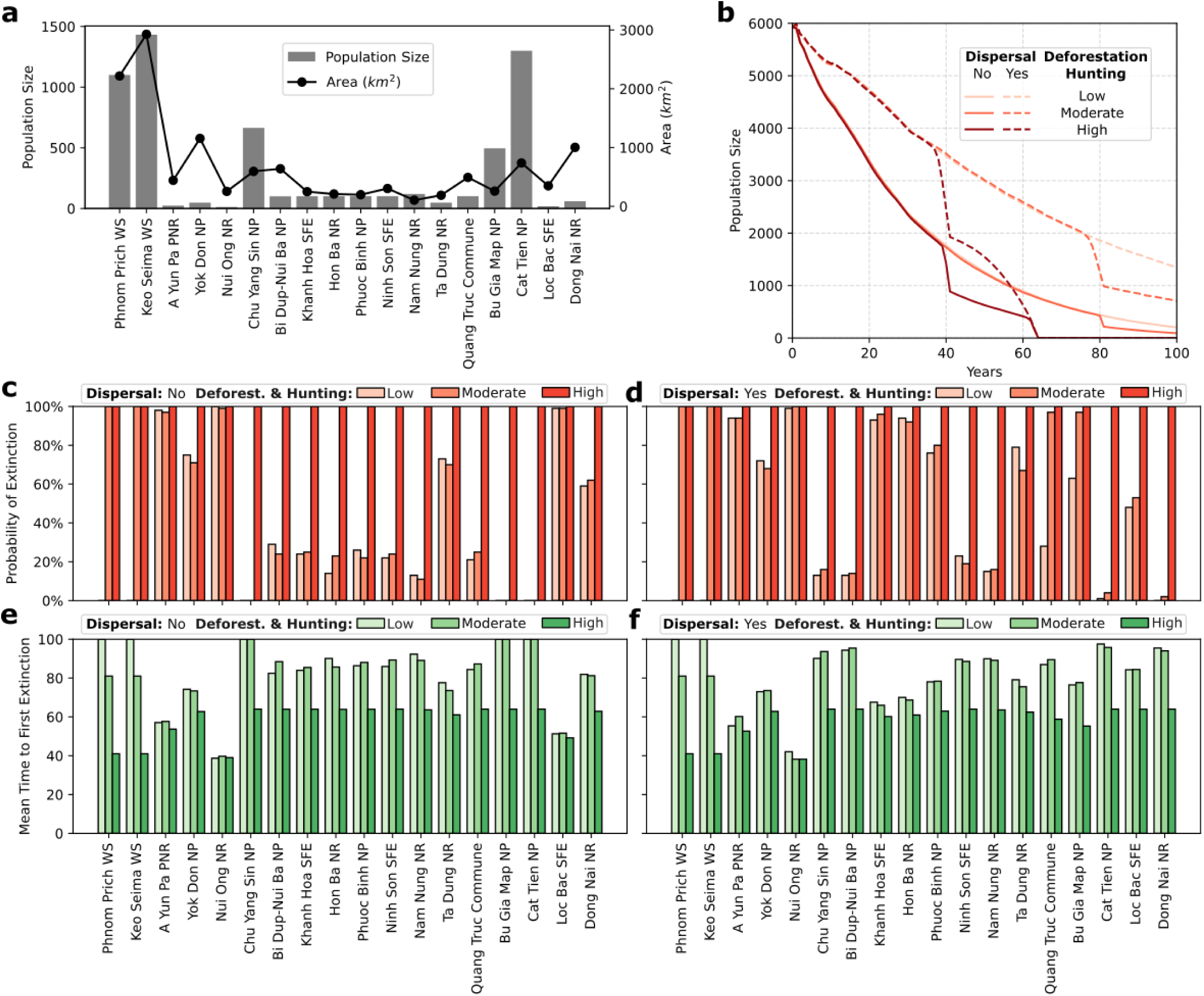
Current state and future population viability of southern yellow-cheeked gibbons. (a) Estimates of current population sizes (grey bars) and areas in km^2^ for sites in Figure 1 that support populations of southern yellow- cheeked gibbons. (b) The changes in the mean meta-population size are predicted from the population viability model. Dispersal of southern yellow-cheeked gibbons between different populations is either ignored (solid lines) or approximated by an ideal free distribution (dashed lines, see Methods). The levels of deforestation and hunting are varied from low (light red), to moderate (red), and high (dark red) levels, as shown in Table 6. (c) Probability of extinction in different sites for different levels of deforestation and hunting in the model with no dispersal. (d) Probability of extinction in different sites for different levels of deforestation and hunting in the model with ideal free dispersal. (e) Mean time to first extinction (in years) for different sites and levels of deforestation and hunting in the model with no dispersal. (f) Mean time to first extinction (in years) for different sites and levels of deforestation and hunting in the model with ideal free dispersal.

**Table 1:**
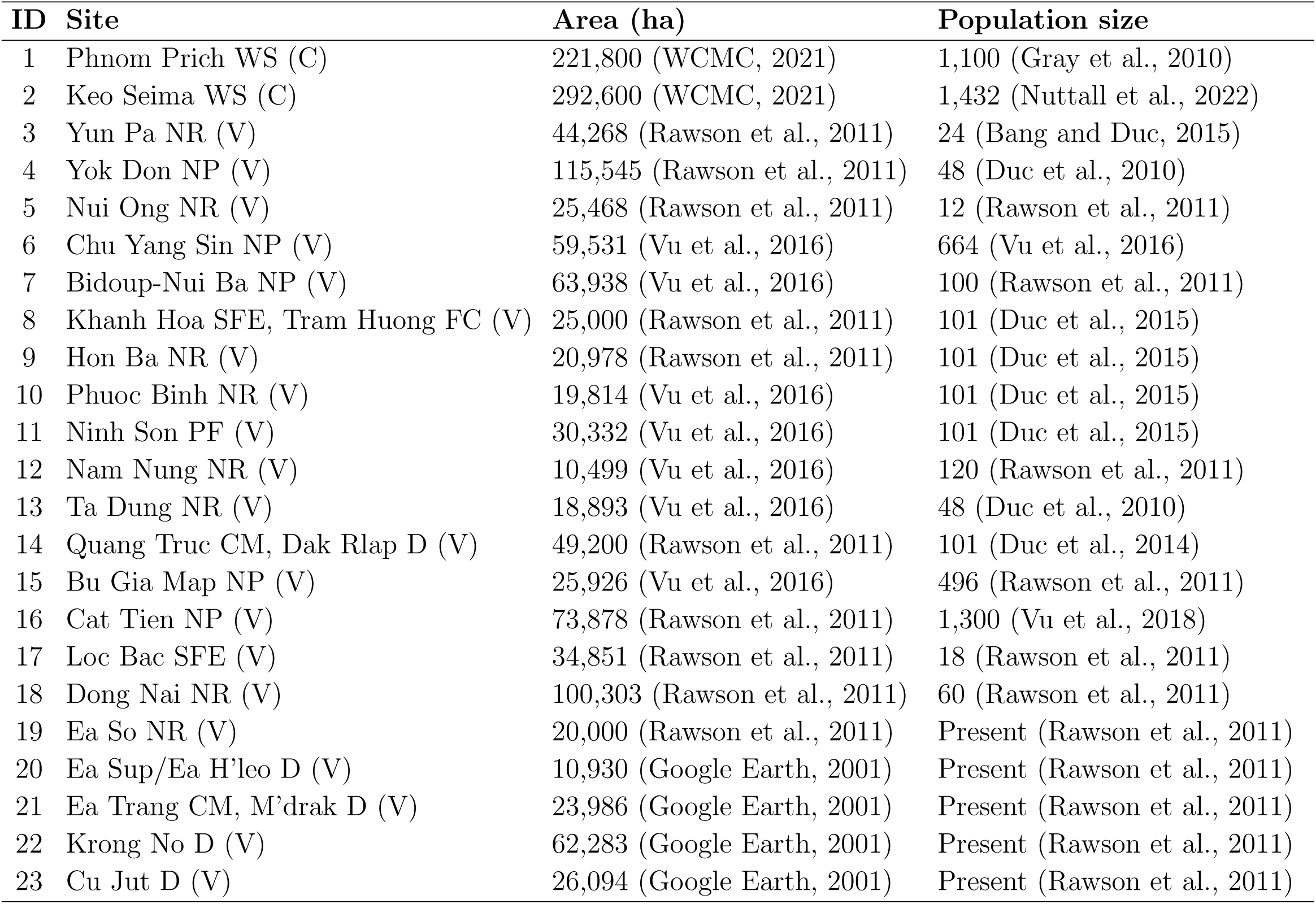
Sites with yellow-cheeked gibbons, their areas and population size. Abbreviations: C=Cambodia, V=Vietnam, WS=Wildlife Sanctuary, NR=Nature Reserve, NP=National Park, PF=Protected Forest, SFE=State Forestry Enterprise, FC=Forest Company, CM=commune, D=district.

| ID | Site | Area (ha) | Population size |
| --- | --- | --- | --- |
| 1 | Phnom Prich WS (C) | 221,800 (WCMC, 2021) | 1,100 (Gray et al., 2010) |
| 2 | Keo Seima WS (C) | 292,600 (WCMC, 2021) | 1,432 (Nuttall et al., 2022) |
| 3 | Yun Pa NR (V) | 44,268 (Rawson et al., 2011) | 24 (Bang and Duc, 2015) |
| 4 | Yok Don NP (V) | 115,545 (Rawson et al., 2011) | 48 (Duc et al., 2010) |
| 5 | Nui Ong NR (V) | 25,468 (Rawson et al., 2011) | 12 (Rawson et al., 2011) |
| 6 | Chu Yang Sin NP (V) | 59,531 (Vu et al., 2016) | 664 (Vu et al., 2016) |
| 7 | Bidoup-Nui Ba NP (V) | 63,938 (Vu et al., 2016) | 100 (Rawson et al., 2011) |
| 8 | Khanh Hoa SFE, Tram Huong FC (V) | 25,000 (Rawson et al., 2011) | 101 (Duc et al., 2015) |
| 9 | Hon Ba NR (V) | 20,978 (Rawson et al., 2011) | 101 (Duc et al., 2015) |
| 10 | Phuoc Binh NR (V) | 19,814 (Vu et al., 2016) | 101 (Duc et al., 2015) |
| 11 | Ninh Son PF (V) | 30,332 (Vu et al., 2016) | 101 (Duc et al., 2015) |
| 12 | Nam Nung NR (V) | 10,499 (Vu et al., 2016) | 120 (Rawson et al., 2011) |
| 13 | Ta Dung NR (V) | 18,893 (Vu et al., 2016) | 48 (Duc et al., 2010) |
| 14 | Quang Truc CM, Dak Rlap D (V) | 49,200 (Rawson et al., 2011) | 101 (Duc et al., 2014) |
| 15 | Bu Gia Map NP (V) | 25,926 (Vu et al., 2016) | 496 (Rawson et al., 2011) |
| 16 | Cat Tien NP (V) | 73,878 (Rawson et al., 2011) | 1,300 (Vu et al., 2018) |
| 17 | Loc Bac SFE (V) | 34,851 (Rawson et al., 2011) | 18 (Rawson et al., 2011) |
| 18 | Dong Nai NR (V) | 100,303 (Rawson et al., 2011) | 60 (Rawson et al., 2011) |
| 19 | Ea So NR (V) | 20,000 (Rawson et al., 2011) | Present (Rawson et al., 2011) |
| 20 | Ea Sup/Ea H'leo D (V) | 10,930 (Google Earth, 2001) | Present (Rawson et al., 2011) |
| 21 | Ea Trang CM, M'drak D (V) | 23,986 (Google Earth, 2001) | Present (Rawson et al., 2011) |
| 22 | Krong No D (V) | 62,283 (Google Earth, 2001) | Present (Rawson et al., 2011) |
| 23 | Cu Jut D (V) | 26,094 (Google Earth, 2001) | Present (Rawson et al., 2011) |

**Table 2:** Estimates of annual deforestation and hunting rates.

| Threat severity | Deforestation |  | Hunting |
| --- | --- | --- | --- |
|  | Cambodia | Vietnam | Female + Infant |
| **Low** | 0.6125% | 0.4% | 0.63% |
| **Moderate** | 1.25% | 0.8% | 1.26% |
| **High** | 2.5% | 1.6% | 2.52% |

### 3.2. Populations face the risk of extinction

Currently, yellow-cheeked gibbons face two main threats: deforestation and hunting (Traeholt et al., 2005; Rawson et al., 2011, 2020; Channa and Gray, 2009), with the estimated rates shown in Table 6. To explore the viability of the fragmented populations in the face of these threats, we used our population viability model (Fig. 2b). Since the dispersal of southern yellow-cheeked gibbons is understudied, we assumed that gibbons either do not disperse or disperse ideally and freely between neighbouring sites in Fig. 1, mimicking two extreme dispersal strategies. Similar, since the estimates of deforestation and hunting are imprecise, we considered three different scenarios corresponding to varied threat severity (Table 6). Crucially, our model predicts a steady decline of the southern yellow-cheeked gibbon meta- population under all considered scenarios (Fig. 2b, Supplementary Table S2). Simulations that include dispersal exhibit a faster decline (Fig. 2b), which can be explained by an increased death rate associated with dispersal. Furthermore, the increase in deforestation and hunting decreases the viability of the metapopulation over the next 100 years, giving rise to sudden declines in population abundance (Fig. 2b). For the estimated values of deforestation and hunting, such a sudden metapopulation collapse appears after around 81 years, while there are two collapses at around 41 and 64 years when the threats reach double the estimated values (Fig. 2b).

To understand this metapopulation dynamics, we need to explore the dynamics of all populations in all fragmented sites (Fig. 2c-f, Supplementary Fig. S3). In particular, we can compare the extinction probabilities (Fig. 2c-d) and the mean times to first extinction (Fig. 2e-f) between different populations and different modelling scenarios. When the threats are high (twice the estimated values), the probability of extinction is almost sure in all considered populations irrespective of dispersal (Fig. 2c-d). Moreover, two major populations in Phnom Prich Wildlife Sanctuary and Keo Seima Wildlife Sanctuary go extinct at around 41 years, while the extinction of other populations follows at around 64 years (Fig. 2e-f), which explains the sudden collapse of the metapopulation. When the threats are moderate (current estimated values), the extinction probability of some populations decreases (Fig. 2c-d). However, the populations in Phnom Prich Wildlife Sanctuary and Keo Seima Wildlife Sanctuary are always predicted to go extinct (Fig. 2c-d) at around 81 years (Fig. 2c-d), giving rise to a sudden collapse of the metapopulation (Fig. 2b). Only when threats are reduced (to 50% of the current estimated values), do the populations in Prich Wildlife Sanctuary and Keo Seima Wildlife Sanctuary survive over the next 100 years (Fig. 2c-d). Moreover, in such cases, the survival of the other major populations in Cat Tien, Chu Yang Sin and Bu Gia Map National Parks is significantly improved (Fig. 2e-f). Interestingly, the major populations with the largest size benefit most from reduced deforestation and hunting, while the remaining smaller populations are susceptible to extinctions irrespective of threat severity (Supplementary Fig. S4).

### 3.3. Reduced hunting improves population viability

While the population viability analysis shows that populations are sensitive to changes in deforestation and hunting, it remains unclear which of these threats drives the extinctions in the model. To understand what drives extinctions, we performed a systematic sensitivity analysis where we independently varied the rates of deforestation and hunting from 50% to 200% of the estimated values (Fig. 3, Supplementary Fig. S5). Since dispersal does not significantly modify the extinction times (Fig. 2b), we decided to focus solely on the model with ideal free dispersal. Importantly, the systematic sensitivity analysis suggests that the extinction probability and the mean time to first extinction are generally more sensitive to hunting than deforestation (Fig. 3). For most sites, the estimated annual hunting rate (1.26% of females with infants) is close to a critical value where the probability of extinction suddenly transitions from low to large. In particular, the model predicts that even a small reduction in hunting in the major sites (Keo Seima and Phnom Prich Wildlife Sanctuaries, Cat Tien, Chu Yan Sin and Bu Gia Map National Parks) can have a profound impact on the population viability in these sites. In contrast, the sites with small estimated populations and areas (such as Nui Ong, Hon Ba and A Yun Pa Nature Reserves, and Khanh Hoa State Forestry Enterprise) exhibit high probabilities of extinction regardless of hunting or deforestation levels, underscoring the importance of preserving large intact habitats that carry large populations. Importantly, the large sites with large populations determine the global extinction risks of the entire metapopulation (Supplementary Fig. S5), implying that the reduction of hunting can significantly improve the survival of the whole metapopulation.

**Figure 3:**
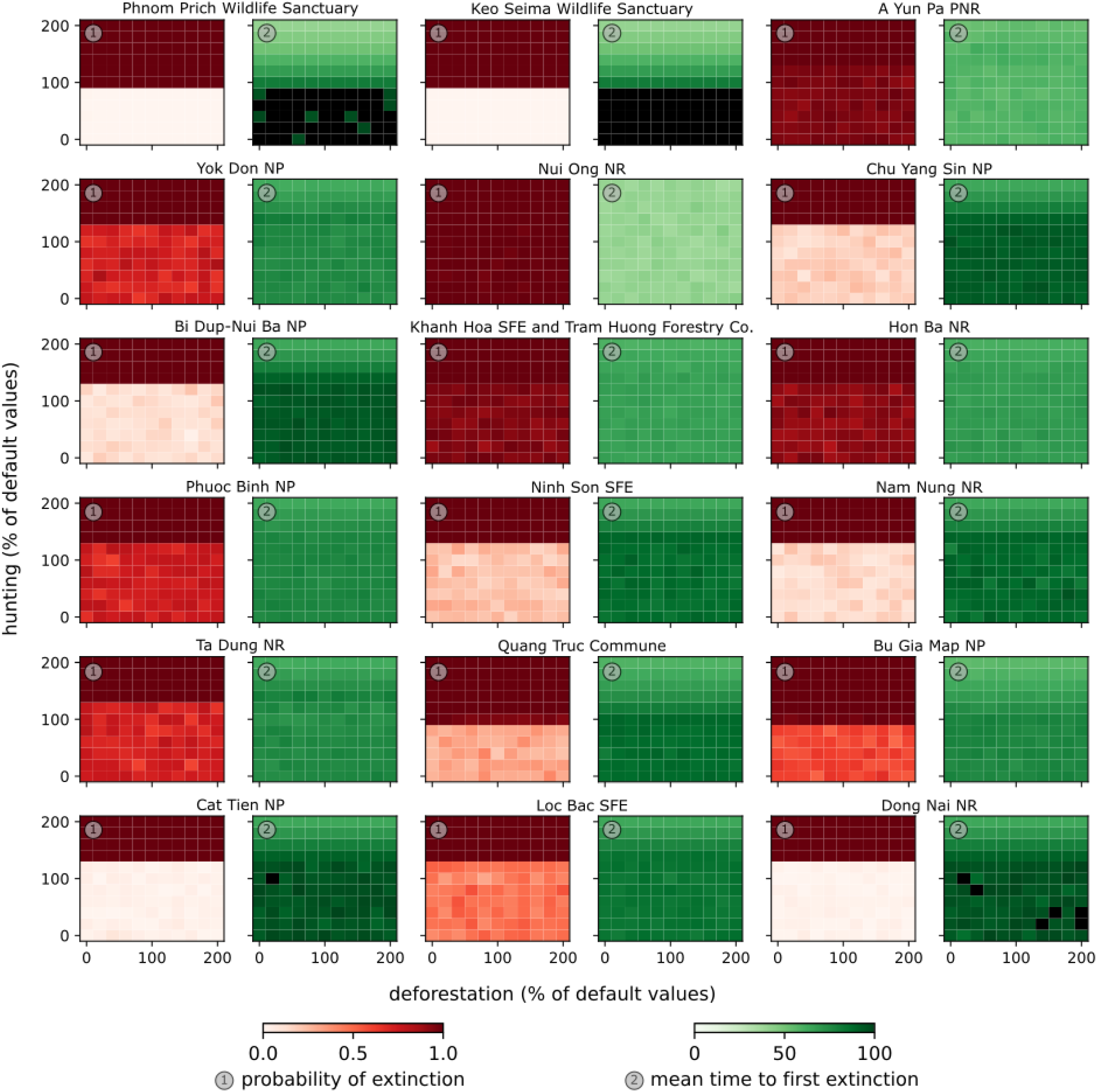
Sensitivity of populations to hunting and deforestation. Probability of extinction (red heatmaps, 1) and mean time to first extinction (measured in years, green heatmaps, 2) are determined for different populations (different plots) and variable levels of deforestation rate (x-axis) and hunting rate (y-axis). The default values of deforestation and hunting rates that correspond to 100% along these axes are given by the moderate values in Table 6. Correspondingly, the low (resp. high) levels of deforestation and hunting in Table 6 correspond to a point with 50% (resp. 200 %) deforestation and hunting rates. The model with the ideal free dispersal is considered.

### 3.4. Conservation of large populations is important

The population viability modelling suggests that the protection of the largest populations plays a key role in conserving the entire metapopulation over the next 100 years (Fig. 2, 3). To confront this prediction with field experience, we conducted online surveys with three experts who conserve the southern yellow-cheeked gibbons in the field. The experts were asked to name and rank the three most important sites for the conservation of yellow-cheeked gibbons in Vietnam and Cambodia. Their responses included the following five sites, ranked in the following order (1) Phnom Prich Wildlife Sanctuary, (2-3) Keo Seima Wildlife Sanctuary and Cat Tien National Park, (4-5) Chu Yan Sin and Bu Gia Map National Parks (Fig. 4a). Crucially, these five sites match exactly with the five most populous sites based on our previous analysis (Fig. 2a), carrying the estimated 84% of the entire metapopulation size. When the experts were asked to provide a reason for their ranking, all three agreed that the population size was the most important factor in their decision. One expert also noted that the contiguity with other populations played an important role in their decision-making: *Seima is connected to Phnom Prich and to Bu Gia Map in Vietnam; Cat Tien is connected to Dong Nai; Chu Yang Sin is connected to a multitude of protected areas/protection forests/extractive reserves*.

**Figure 4:**
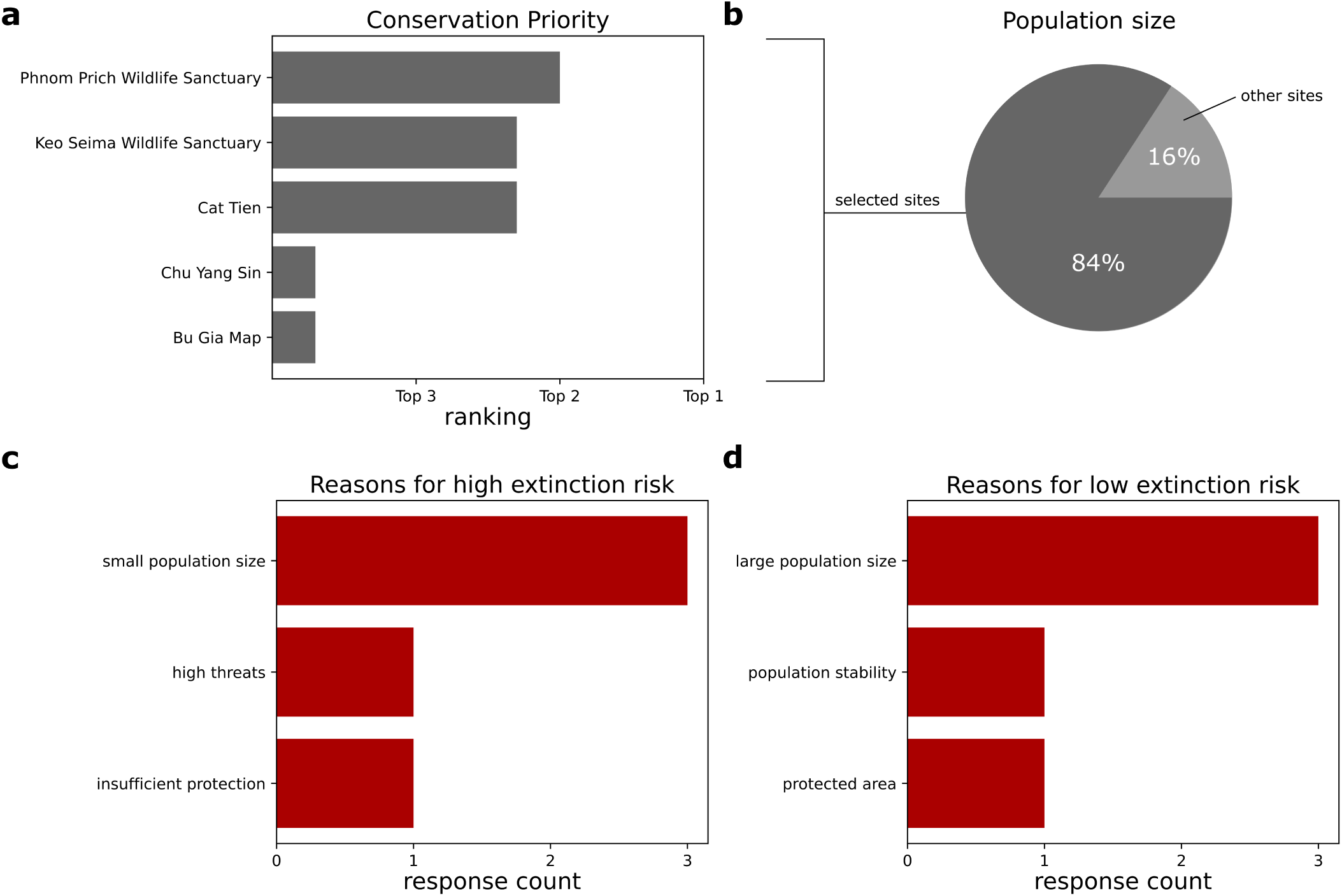
Expert assessment (N=3) of priority areas for conservation and extinction risks. (a) Conservation priority of areas selected by experts. Experts were asked to name the top three areas based on their judgement of conservation priority for southern yellow-cheeked gibbons. The average ranking of selected areas is plotted (see Methods). (b) Total population size of areas selected by experts in part (a). All experts state a large population size as the main reason for their choice of sites in part (a). (c) Reasons for high extinction risk mentioned by the experts. The bar plot shows the number of experts who mentioned a given reason for high extinction risk. (d) Reasons for low extinction risk mentioned by the experts. The bar plot shows the number of experts who mentioned a given reason for low extinction risk.

To elaborate on their reasoning further, the experts were asked to name the sites with the lowest and highest extinction risk for southern yellow-cheeked gibbons and to explain their reasoning. All three experts agreed that population size is the key driver of local extinctions, while one expert additionally provided site protection as an important factor and another expert additionally provided threats and population stability as important factors (Fig. 4c,d). The sites identified by experts as the sites with the highest extinction risk were Bi Dup-Nui Ba and Yok Don National Parks, which are estimated to contain 100 and 48 individuals (Table 6). In contrast, the sites identified as the sites with the lowest extinction risk were Phnom Prich and Keo Seima Wildlife Sanctuaries, which are estimated to contain 1100 and 1432 individuals (Table 6).

### 3.5. Conservation is a complex socio-ecological enterprise: the case study of Cat Tien

While the first part of the survey was designed to confront the population viability model with field experience, its latter part focused on the socio-ecological contexts of gibbon conservation. In this part, the experts were asked to choose a site they were most familiar with. Two of the three experts participated in this part and chose Cat Tien National Park, which we will use to illustrate the complex socio-ecological contexts of conservation.

In the first task, the experts were asked to name stakeholders that affect the current state of yellow-cheeked gibbons in Cat Tien and describe their roles. Based on their answers, we identified the following stakeholders and their relationship to southern yellow-cheeked gibbons (Fig. 5a):

- National Assembly of Vietnam: in charge of protection laws, which are not sufficient;
- Vietnam Administration of Forestry: provides funding and resources;
- Cat Tien National Park management board: manages the protected area;
- Rangers and guides: protect against poachers; explain the necessity for the protection and for the national park;
- Tourists: provide funding; are witnesses (“eyes in the forest”); but also feed the wildlife; and leave garbage in the forest;
- Poachers: kill or capture gibbons; still able to enter the site despite all the good efforts the rangers make;
- Local people: extract and trade forest products, which may affect gibbon habitat; villages at the park edge contribute to deforestation by burning down trees.

**Figure 5:**
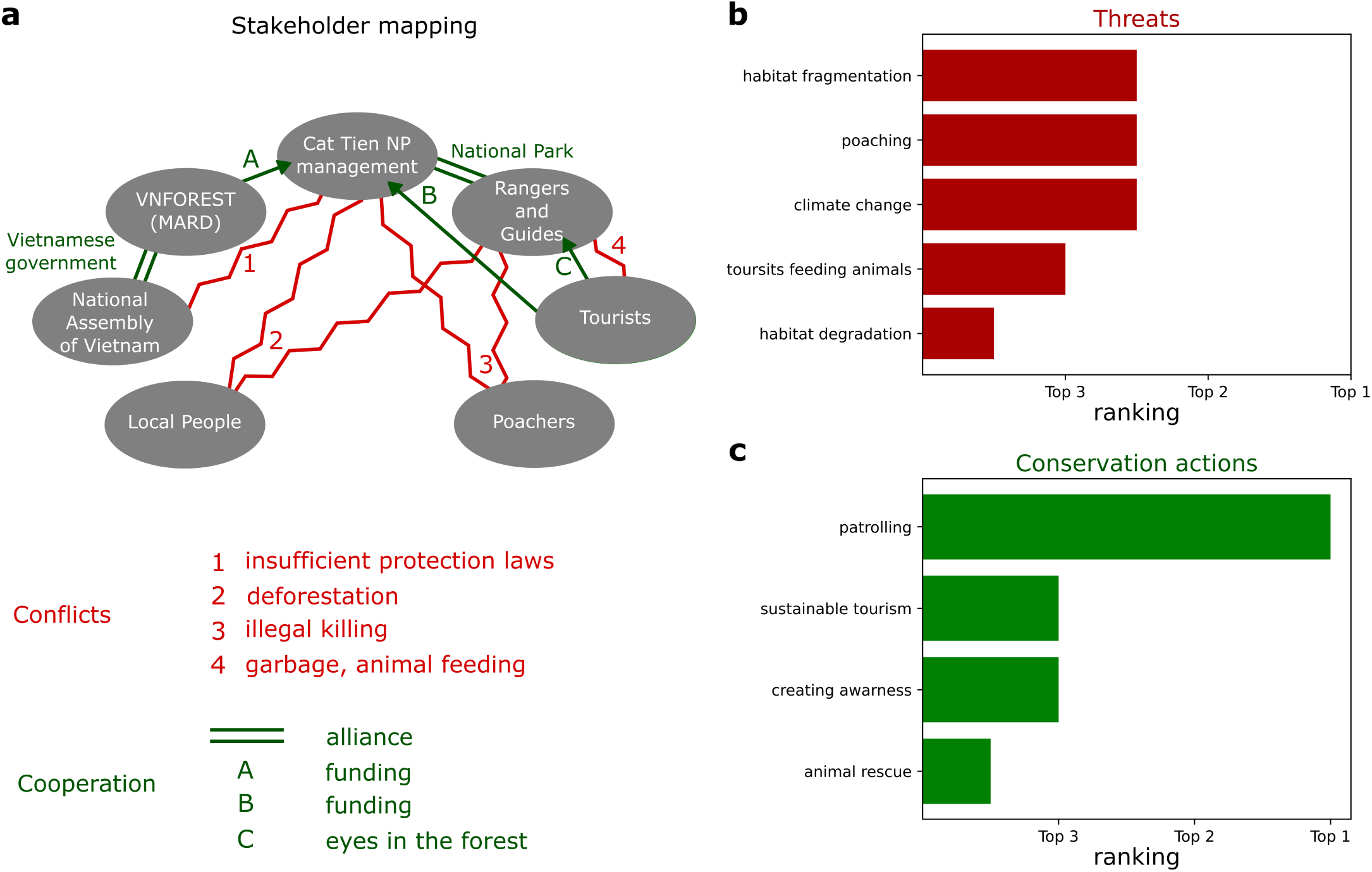
Expert assessment (N=3) of the conservation of southern yellow-cheeked gibbons in Cat Tien National Park. (a) Stakeholder mapping for Cat Tien National Park. Based on the expert surveys, stakeholders affecting the conservation of southern yellow-cheeked gibbons were identified (ellipses), including potential cooperative interactions (green straight lines) and conflicts (red zig-zag lines). Abbreviations: VNFOREST = Vietnam Administration of Forestry, MARD = Ministry of Agriculture and Rural Development, NP = National Park. (b) Threats posed to southern yellow-cheeked gibbons selected by the experts (red). Experts were asked to name the top three threats to southern yellow-cheeked gibbons based on their judgement. The average ranking of selected threats is plotted. (c) Types of conservation actions proposed by the experts (green). Experts were asked to name the top three conservation actions for the southern yellow-cheeked gibbons based on their judgement. The average ranking of selected conservation actions is plotted.

These relationships imply several conflicts and collaborations between the stakeholders, revealing how complex human social dynamics affect the conservation of southern yellow-cheeked gibbons. Importantly, the management board, rangers and guides of the national park form a key alliance that co- operates to conserve the southern yellow-cheeked gibbons. While this alliance receives funding from the Vietnamese government and tourists, their relationship with these stakeholders is ambivalent. For example, the Vietnamese government includes the National Assembly of Vietnam and the Vietnam Administration of Forestry, which is a government agency under the Ministry of Agriculture and Rural Development. While the Vietnam Administration of Forestry cooperates on conservation by providing funding and resources, the National Assembly of Vietnam creates a conflict due to insufficient protection laws (Fig. 5a). Similarly, the relationship between the national park and tourists is multifaceted. On one hand, the tourists provide the national park with funding and help the rangers as eyewitnesses in the forest. On the other hand, they feed the animals and leave garbage in the park. The situation with garbage is particularly complicated. Unfortunately, there is no garbage collection for the park lodge, which the national park resolves by burning the garbage directly in the tourist part of the park. Such a solution increases the carbon footprint, attracts Macaques, and undermines the credibility of the values and efforts of the national park. Finally, the relationship between the national park, poachers and local people is complicated. The local people cause deforestation by burning trees and extracting and trading forest products, degrading the gibbon habitat. Moreover, despite the good efforts of rangers, the poachers can enter the site and illegally hunt the southern yellow-cheeked gibbons. In summary, the stakeholder mapping uncovers intricate social dynamics among stakeholders, including cooperative interactions (alliances, funding support, park surveillance support) and conflicting interactions (inadequate legal support, deforestation, illegal hunting, improper waste disposal and animal feeding practices).

To further elaborate on the relationship between humans and southern yellow-cheeked gibbons, the experts were asked to name and rank the three biggest threats to the gibbons in Cat Tien. The first expert ranked the threats from most severe to least severe as: (1) poaching, (2) inappropriate tourist behaviour (animal feeding) and (3) climate change. This expert further noted that the feeding by tourists disrupts the natural behaviour of gibbons. The second expert ranked the threats from most severe to least severe as: (1) habitat fragmentation, (2) climate change and (3) habitat degradation. This expert explained that habitat fragmentation promotes the isolation of populations, which is a severe threat due to the persistence of lacking connectivity between adjacent forests. Moreover, according to this expert, climate change poses a threat over the medium term of the next five gibbon generations with possibly highly detrimental consequences for the gibbon habitat. Finally, the expert concluded that logging and non-timber forest products extraction cause habitat degradation, even though practically all high-value timber in accessible areas has already been removed. In summary, the three major threats identified by the experts are habitat fragmentation, poaching and climate change, and these threats are followed by animal feeding and habitat degradation (Figure 5b).

To identify effective solutions to these threats, the experts were asked to provide the three most effective conservation actions for the southern yellow-cheeked gibbons in Cat Tien. The first expert identified two actions in order of effectiveness, from most to least: (1) deploying rangers and (2) raising awareness. The second expert identified three actions in order of effectiveness, from most to least, making the following comments: (1) effective patrolling, (2) sustainable tourism (“*puts a value on the park and the gibbons, pushing conservation up the political agenda*”) and (3) animal rescue centre (“*animals that are wounded/sick can get treatment at the Rescue Center and can be released back if possible*”). Regarding effective patrolling, this expert noted: *“Ultimately, it is the encouragement of more sustainable livelihoods in the region which will decide the fate of the population, but that is beyond the scale/remit of what conservation agencies can influence. Patrolling is just a stop-gap. It seems that some rangers do their utmost best, but their salary is very low, so many go for another job or even become or help poachers. Rangers and guides really do their best to create awareness but have little means. In the park, they use diesel cars for safari cars so there is room for eco-friendly electric cars.”* In summary, both experts mentioned the patrolling of rangers as the first and most effective conservation action (Figure 5c).

## 4. Discussion

In this work, we combined population viability modelling and expert surveys to examine the conservation of southern yellow-cheeked gibbons in Vietnam and Cambodia. While the population viability modelling can guide conservation strategies by predicting future population trends (Fig. 2, 3;Souĺe, 1985; Stark et al., 2012; Marshall et al., 2008; Tunhikorn et al., 1994; Molur et al., 2005; Fan et al., 2013; Bryant, 2014; Smith et al., 2017), this method cannot incorporate any socio-ecological complexity of the gibbon conservation and requires integration with a social study to address this complexity. By incorporating the expert surveys and stakeholder mapping (Fig. 4, 5), our work contributes to the field of ethnoprimatology (Fuentes, 2010, 2012; Dore et al., 2017; Cheyne et al., 2023), that recognizes the importance of both biological and social aspects of nature conservation. Our study opens a potential for further research in this socio-ecological area that would involve more stakeholders to resolve some of their conflicts (Redpath et al., 2013). In particular, our stakeholder mapping uncovers a complex relationship between the protected areas, hunters and local people (Fig. 5) and further surveys with the local communities could foster mutual understanding and empower the voice of local communities in the spirit of decolonizing conservation (Adams and Mulligan, 2012).

In addition to further social studies, there is a necessity for further biological monitoring of the populations. While the large populations in Phnom Phrich Wildlife Sanctuary (Channa and Gray, 2009; Gray et al., 2010), Keo Seima Wildlife Sanctuary (Rawson et al., 2009; Pollard et al., 2007), Cat Tien National Park (Kenyon, 2008; Kenyon et al., 2011) and Chu Yang Sin National Park (Vu et al., 2016) have been monitored, there is a need for improved monitoring. Firstly, new technologies, such as GPS collars (Kenyon et al., 2015) and drones equipped with cameras (Wearn et al., 2023), can significantly improve the current estimation of individual counts that have traditionally relied on indirect inference from vocalizations (Wearn et al., 2024). Since the population sizes are a key input of the population viability model (Fig. 2), these new survey techniques could improve the precision of the model. Secondly, there is a need to conduct population surveys in understudied areas that could potentially carry significant populations of southern yellow-cheeked gibbons. Previous habitat suitability analysis models suggest that evergreen forests in Dak Lak, Dak Nong, Lam Dong, Dong Nai, and Binh Phuoc provinces in Vietnam and in Mundulkiri in Cambodia are suitable for this gibbon species (Vu et al., 2018; Nhung et al., 2021). The community-based monitoring in the Da Te forest (Dang and Vinh, 2011) and the survey in Ta Dung Nature Reserve (Duc et al., 2010) provide great examples of survey efforts in the understudied areas. In particular, it seems important to understand the populations in the regions that connect the major Cambodian population in Keo Seima Wildlife Sanctuary to the major Vietnamese populations in Bu Gia Map and Cat Tien National Parks (Fig. 1), such as Vinh An, Ma Da, Hieu Liem, Da Teh, Loc Bac and Bao Lam State Forestry Enterprises (Rawson et al., 2011).

The distribution of the southern yellow-cheeked gibbons between Cambodia and Vietnam (Fig. 1) necessitates a transboundary conservation approach that would ensure habitat connectivity (Mason et al., 2020). Importantly, the Greater Mekong Subregion Biodiversity Conservation Corridors Initiative (Greater Mekong Subregion Core Environment Program, 2008) was established with the goal to promote contiguity of transboundary populations in South East Asia, including the contiguity of Keo Seima Wildlife Sanctuary in Cambodia and Bu Gia Map National Park in Vietnam (Fig. 1). While the transboundary efforts of this project continue in some regions of South East Asia, such as for the Cao- vit gibbon (*Nomascus nasutus*) on the border between Vietnam and China (Wang et al., 2021; Ma et al., 2020), the southern yellow-cheeked gibbons would benefit from further transboundary collaborations between Vietnam and Cambodia. Such a transboundary collaboration between Vietnam and Cambodia would fulfill the goals of the Convention on Biological Diversity (CBD, 1992) and the Convention on International Trade in Endangered Species of Wild Fauna and Flora (CITES, 1973) that both countries signed.

In addition to the international conventions, both Vietnam and Cambodia have their own legislation that protects southern yellow-cheeked gibbons. In Cambodia, the Cambodian Forestry Law (2002) and the Sub-decree No. 53 (2006) forbid the hunting, trapping, possession, transport, and trade of any endangered species, including southern yellow-cheeked gibbons. In Vietnam, these activities are forbidden by Decrees No. 82/2006/ND-CP, 32/2006/ND-CP, 160/2013/ND-CP. Furthermore, the habitat of southern yellow-cheeked gibbons is protected in designated protected areas by Cambodian Law on Protected Areas (2008) and Vietnamese Law on Biodiversity (2008) and Decree No. 109/2003/ND-CP. However, our expert surveys indicate that there is a need to improve these laws (Fig. 5), for example, by including protecting biodiversity corridors between different protected areas, reducing logging in key habitats, improving the enforcement of these laws, or creating animal welfare legislation (Ibbett et al., 2021; Traeholt et al., 2005; Rawson et al., 2011).

While the local laws in Vietnam and Cambodia should, in principle, protect the southern yellow-cheeked gibbons from hunting, hunting is still common (Ibbett et al., 2021) and constitutes the key threat to the gibbons (Fig. 3, 5; Rawson et al., 2011; 2005). Hunting is deeply embedded in the culture and includes hunting for wild meat and for illegal pet trade. While 84% households in Keo Seima consume wild meat (Ibbett et al., 2021), southern yellow-cheeked gibbons are primarily hunted for pet trade (Rawson et al., 2011; Geissmann, 2007). Captive gibbons are common in hotels and private zoos in southern Vietnam and it is likely that these are sourced from the wild (Rawson et al., 2011), with increasing online market complicating the enforcement of laws that forbid hunting (Smith et al., 2017). Moreover, if such gibbons are saved, they are commonly severly starved and require medical help, including complex surgeries (Duy et al., 2010).

In addition to reducing hunting, there is a need to protect the habitat of gibbons that is increasingly fragmented and reduced (Fig. 1, 5; Rawson et al., 2011). The agricultural encroachment and logging reduce habitat connectivity, allow hunters to access the habitat, and possibly isolate gibbon groups that become prone to inbreeding (Rawson et al., 2011). Preserving gibbon habitats is particularly important in the face of climate change, that was not explicitly included in our model but has been studied extensively (Yang et al., 2023; Vu et al., 2018; Nhung et al., 2021). These studies predict that multiple gibbon species risk the loss of up to 46% of their habitat by 2050 due to climate change (Yang et al., 2023), with southern yellow-cheeked gibbons risking the loss up to 66% of their habitat by 2050 (Vu et al., 2018) or more optimistic 40% by 2060 (Nhung et al., 2021).

Such negative prognoses require us to preserve some populations in captivity. Captive populations are not only important as genetic banks, but also for gibbons rescued from pet trade to heal and for us to better understand them. The rescued individuals that heal can be reintroduced into the wild (Kenyon et al., 2015), while the captive individuals in zoos have provided us with the understanding of the gibbon reproductive cycle (Fan et al., 2021) which we used in our model. Moreover, the study of gibbon vocalizations in captivity (Hradec et al., 2017, 2021b,a) has the potential to be used in improving the monitoring methods of wild populations that are heavily dependent on recording vocalizations.

Overall, the southern yellow-cheeked gibbons are a unique gibbon species with importance for entire ecosystems (Hai et al., 2018) and deserve dedicated conservation effort. Our study shows that hunting, habitat fragmentation and climate change threaten the survival of this unique species (Fig. 5b). However, effective implementation of conservation actions (Fig. 5c) can reduce these threats and save the southern yellow-cheeked gibbons from extinction (Fig. 2c,d; Fig. 3).

## Acknowledgements

We thank Chloe Montes Strevens for helpful comments on an early version of this work. P.P. was supported by a research scholarship from the School of Geography and the Environment, University of Oxford and a conference and fieldwork funding from Green Templeton College, Oxford. V.P. was supported by the University of Oxford Mathematical Institute Scholarship.

## 5. Author contributions

**Pavla Piskovska**: Conceptualization, Data curation, Formal analysis, Funding acquisition, Investigation, Methodology, Project administration, Software, Validation, Visualization, Writing – original draft, Writing – review and editing; **Vit Piskovsky**: Software, Visualization, Writing – original draft, Writing – review and editing; **Susan Cheyne**: Conceptualization, Supervision, Writing – review and editing.

## References

1. Adams, W.B., Mulligan, M., 2012. Decolonizing nature: strategies for conservation in a post-colonial era. Routledge.

2. Almeida-Rocha, J.M.d., Peres, C.A., Oliveira, L.C., 2017. Primate responses to anthropogenic habitat disturbance: A pantropical meta-analysis. Biol. Conserv. 215, 30–38.

3. Azhar, B., Lindenmayer, D., Wood, J., Fischer, J., Manning, A., McElhinny, C., Zakaria, M., 2013. Contribution of illegal hunting, culling of pest species, road accidents and feral dogs to biodiversity loss in established oil-palm landscapes. Wildl. Res. 40, 1.

4. Bang, T.V., Duc, H.M., 2015. A yun pa propose natural reserve: Initial data on primate fauna, in: Proceeding of the 6th National Scientific Conference on Ecology and Biological Resources, Hanoi.

5. Barca, B., Vincent, C., Soeung, K., Nuttall, M., Hobson, K., 2016. Multi-female group in the southern- most species of nomascus: field observations in eastern Cambodia reveal multiple breeding females in a single group of southern yellow-cheeked crested gibbon *nomascus gabriellae*. Asian Primates Journal 6, 15–19.

6. Benitez-Lopez, A., Alkemade, R., Schipper, A.M., others, 2017. The impact of hunting on tropical mammal and bird populations. Science .

7. Brockelman, W.Y., Reichard, U., Treesucon, U., Raemaekers, J.J., 1998. Dispersal, pair formation and social structure in gibbons (*Hylobates lar* ). Behav. Ecol. Sociobiol. 42, 329–339.

8. Brodie, J.F., Giordano, A.J., Zipkin, E.F., Bernard, H., Mohd-Azlan, J., Ambu, L., 2015. Correlation and persistence of hunting and logging impacts on tropical rainforest mammals: Logging, hunting, and mammal diversity. Conserv. Biol. 29, 110–121.

9. Bryant, J.V., 2014. Developing a conservation evidence-base for the Critically Endangered Hainan gibbon (’Nomascus hainanus’). Ph.D. thesis. UCL.

10. Caldecott, J.O., Miles, L., 2005. World Atlas of Great Apes and Their Conservation. University of California Press.

11. Cambodian Ministry of Environment, UN Environment Programme WCMC, The Ramsar Convention Secretariat, 2023. Dataset: Natural protected areas in Cambodia (1993-2023). URL: https://data.opendevelopment{Cxambodia.net/dataset/protectedareas.

12. CBD, 1992. Convention on Biological Diversity. Signed in Rio de Janeiro, Brazil. Available at:https://www.cbd.int/.

13. Channa, P., Gray, T., 2009. The status and habitat of yellow-cheeked crested gibbon nomascus gabriellae in phnom prich wildlife sanctuary, mondulkiri. Phnom Penh: WWF Greater Mekong-Cambodia .

14. Chatterjee, H.J., 2006.Phylogeny and biogeography of gibbons: A dispersal-vicariance analysis. Int.J. Primatol. 27, 699–712.

15. Cheyne, S.M., Thompson, C., Fan, P.F., Chatterjee, H.J., 2023. Gibbon Conservation in the Anthropocene. Cambridge University Press.

16. CITES, 1973. Convention on International Trade in Endangered Species of Wild Fauna and Flora. Signed in Washington, D.C. Available at: https://cites.org/.

17. Collard, R.C., Dempsey, J., Sundberg, J., 2015. A manifesto for abundant futures. Ann. Assoc. Am. Geogr. 105, 322–330.

18. Cowlishaw, G., Dunbar, R.I.M., 2021. Primate Conservation Biology. University of Chicago Press

19. Dang, N.X., Vinh, N.X., 2011. Community-based monitoring of southern yellow-cheeked gibbon *Nomascus gabriellae* in Da Te forest, Lam Dong province, central Vietnam. Vietnamese Journal of Primatology 5.

20. Didham, R.K., Tylianakis, J.M., Gemmell, N.J., Rand, T.A., Ewers, R.M., 2007. Interactive effects of habitat modification and species invasion on native species decline. Trends Ecol. Evol. 22, 489–496.

21. Dore, K.M., Riley, E.P., Fuentes, A., 2017. Ethnoprimatology. Cambridge University Press.

22. Duc, H.M., Bang, T., Covert, H., 2015. Assessment of conservation status and strengthening conservation of yellow-cheeked crested gibbon and other primates in southeastern slope of the Da Lat Plateau, Vietnam, in: Ho Chi Minh Final report to Southern Institute of Ecology and US Fish and Wildlife Service.

23. Duc, H.M., Bang, T.V., Tinh, M.X., Thang, N.D., 2014. Final Report on Fauna in Quang Truc Commune, Tuy Duc District, Dak Nong Province with the emphasis on population assessment of yellow-cheeked gibbon. Technical Report. Southern Institute of Ecology and Wildlife Conservation Society – Vietnam Programme.

24. Duc, H.M., Van Bang, T., Long, V., 2010. Population status of the yellow-cheeked crested gibbon (nomascus gabriellae) in ta dung nature reserve, dak nong province, vietnam. Fauna & Flora International and Conservation International, Hanoi, Vietnam 22.

25. Dunay, E., Apakupakul, K., Leard, S., Palmer, J.L., Deem, S.L., 2018. Pathogen transmission from humans to great apes is a growing threat to primate conservation. Ecohealth 15, 148–162.

26. Duy, T.P., Hoang, T.H., van den Bos, F., Kenyon, M., 2010. Successful cataract removal, and lens replacement on a rescued yellow-cheeked gibbon (*nomascus gabriellae*). Vietnamese Journal of Primatology 4, 69–74.

27. ESRI, 2021. Dataset: World topographic map. URL: https://services.arcgisonline.com/ArcGIS/rest/services/World_Topo_Map/MapServer.

28. Fan, P., Bartlett, T.Q., 2017. Overlooked small apes need more attention! Am. J. Primatol. 79.

29. Fan, P., He, X., Yang, Y., Liu, X., Zhang, H., Yuan, L., Chen, W., Liu, D., Fan, P., 2021. Reproductive parameters of captive female northern white-cheeked (*nomascus leucogenys*) and yellow-cheeked (*nomascus gabriellae*) gibbons. Int. J. Primatol. 42, 49–63.

30. Fan, P.F., Ren, G.P., Wang, W., Scott, M.B., Ma, C.Y., Fei, H.L., Wang, L., Xiao, W., Zhu, J.G., 2013. Habitat evaluation and population viability analysis of the last population of cao vit gibbon (nomascus nasutus): Implications for conservation. Biol. Conserv. 161, 39–47.

31. Fuentes, A., 2010. Naturalcultural encounters in bali: Monkeys, temples, tourists, and ethnoprimatology. Cultural Anthropology: Journal of the Society for Cultural Anthropology 25, 600–624.

32. Fuentes, A., 2012. Ethnoprimatology and the anthropology of the human-primate interface. Annual Review of Anthropology 41, 101–117.

33. Gates, J.F., 1996. Habitat alteration, hunting and the conservation of folivorous primates in african forests. Aust. J. Ecol. 21, 1–9.

34. Geissmann, T., 2000. Vietnam primate conservation status review 2000, part 1 : gibbons. Fauna and Flora International, Indochina Programme, Hanoi, 2000 .

35. Geissmann, T., 2007. Status reassessment of the gibbons: results of the asian primate red list workshop 2006. Gibbon Journal 3, 5–15.

36. Google Earth, 2001. Google Earth. URL: https://earth.google.com/web/.

37. Gray, T.N.E., Phan, C., Long, B., 2010. Modelling species distribution at multiple spatial scales: gibbon habitat preferences in a fragmented landscape. Anim. Conserv. 13, 324–332.

38. Greater Mekong Subregion Core Environment Program, 2008. Biodiversity Conservation Corridors Initiative: Pilot Site Implementation Status Report 2007. Technical Report. Asian Development Bank.

39. Haddad, N.M., Brudvig, L.A., Clobert, J., Davies, K.F., Gonzalez, A., Holt, R.D., Lovejoy, T.E., Sexton, J.O., Austin, M.P., Collins, C.D., Cook, W.M., Damschen, E.I., Ewers, R.M., Foster, B.L., Jenkins, C.N., King, A.J., Laurance, W.F., Levey, D.J., Margules, C.R., Melbourne, B.A., Nicholls, A.O., Orrock, J.L., Song, D.X., Townshend, J.R., 2015. Habitat fragmentation and its lasting impact on Earth’s ecosystems. Sci Adv 1, e1500052.

40. Hai, B.T., Chen, J., McConkey, K.R., Dayananda, S.K., 2018. Gibbons *nomascus gabriellae* provide key seed dispersal for the Pacific walnut (*dracontomelon dao*), in Asia’s lowland tropical forest. Acta Oecol. 88, 71–79.

41. Hansen, S., Roets, F., Seymour, C.L., van Veen, F.J.F., Pryke, J.S., Théebault, E.S., 2018. Alien plants have greater impact than habitat fragmentation on native insect flower visitation networks. Diversity and Distributions 24, 58–68.

42. Hanski, I., 2011. Habitat loss, the dynamics of biodiversity, and a perspective on conservation. Ambio 40, 248–255.

43. Harfoot, M.B.J., Johnston, A., Balmford, A., Burgess, N.D., Butchart, S.H.M., Dias, M.P., Hazin, C., Hilton-Taylor, C., Hoffmann, M., Isaac, N.J.B., Iversen, L.L., Outhwaite, C.L., Visconti, P., Geldmann, J., 2021. Using the IUCN red list to map threats to terrestrial vertebrates at global scale. Nat Ecol Evol 5, 1510–1519.

44. Hien, N.T.T., Binh, N.T., 2024. An overview of the yellow-checked gibbon (nomascus gabriellae thomas, 1909). Journal of Thu Dau Mot University , 207–213.

45. Hradec, M., Illmann, G., Bartšs, L., Bolechová, P., 2021a. The transition from the female-like great calls to male calls during ontogeny in southern yellow-cheeked gibbon males *nomascus gabriellae*. Sci. Rep. 11, 22040.

46. Hradec, M., Illmann, G., Bolechová, P., 2021b. A first report of separation calls in southern yellow-cheeked gibbons (*Nomascus gabriellae*) in captivity. Primates 62, 5–10.

47. Hradec, M., Linhart, P., Bartšs, L., Bolechová, P., 2017. The traits of the great calls in the juvenile and adolescent gibbon males *nomascus gabriellae*. PLoS One 12, e0173959.

48. Ibbett, H., Keane, A., Dobson, A.D.M., Griffin, O., Travers, H., Milner-Gulland, E.J., 2021. Estimating hunting prevalence and reliance on wild meat in cambodia’s eastern plains. Oryx 55, 878–888.

49. IUCN, 2023. IUCN Red List of Threatened Species. URL: https://www.iucnredlist.org. accessed: January 21, 2023.

50. Junker, J., Petrovan, S.O., Arroyo-Rodŕıguez, V., Boonratana, R., Byler, D., Chapman, C.A., Chetry, D., Cheyne, S.M., Cornejo, F.M., Cortés-Ortiz, L., Cowlishaw, G., Christie, A.P., Crockford, C., de la Torre, S., de Melo, F.R., Fan, P., Grueter, C.C., Guzmán-Caro, D.C., Heymann, E.W., Herbinger, I., Hoang, M.D., Horwich, R.H., Humle, T., Ikemeh, R.A., Imong, I.S., Jerusalinsky, L., Johnson, S.E., Kappeler, P.M., Kierulff, M.C.M., Końe, I., Kormos, R., Le, K.Q., Li, B., Marshall, A.J., Meijaard, E., Mittermeier, R.A., Muroyama, Y., Neugebauer, E., Orth, L., Palacios, E., Papworth, S.K., Plumptre, A.J., Rawson, B.M., Refisch, J., Ratsimbazafy, J., Roos, C., Setchell, J.M., Smith, R.K., Sop, T., Schwitzer, C., Slater, K., Strum, S.C., Sutherland, W.J., Talebi, M., Wallis, J., Wich, S., Williamson, E.A., Wittig, R.M., Kühl, H.S., 2021. Corrigendum: A severe lack of evidence limits effective conservation of the world’s primates. Bioscience 71, 105.

51. Kenyon, M., Roos, C., Binh, V.T., Chivers, D., 2011. Extrapair paternity in golden-cheeked gibbons (nomascus gabriellae) in the secondary lowland forest of cat tien national park, Vietnam. Folia Primatol. 82, 154–164.

52. Kenyon, M., Streicher, U., Pei, K.J.C., Cronin, A., van Dien, N., van Mui, T., van Hien, L., 2015. Experiences using VHF and VHF/GPS-GSM radio-transmitters on released southern yellow-cheeked gibbons (*nomascus gabriellae*) in South Vietnam. Vietnamese Journal of Primatology 2, 15–27.

53. Kenyon, M.A., 2008. Ecology of the golden-cheeked gibbon (*nomascus gabriellae*) in Cat Tien National Park, Vietnam. Ph.D. thesis. University of Cambridge.

54. Krivan, V., Cressman, R., Schneider, C., 2008. The ideal free distribution: a review and synthesis of the game-theoretic perspective. Theor. Popul. Biol. 73, 403–425.

55. Lacy, R., Pollak, J., 2021. Vortex: A stochastic simulation of the extinction process.

56. Lappan, S., Ruppert, N., 2019. Primate research and conservation in Malaysia. CAB Rev. Perspect. Agric. Vet. Sci. Nutr. Nat. Resour. 2019, 1–10.

57. Lewis, S.L., Maslin, M.A., 2015. Defining the anthropocene. Nature 519, 171–180.

58. Ma, C.Y., Hoang, T.D., Nguyen, V.T., Le, T.D., Le, V.D., Le, H.O., Yang, J., Zhang, Z.J., Fan, P.F., 2020. Transboundary conservation of the last remaining population of the Cao Vit gibbon *Nomascus Nasutus*. Oryx: The Journal of the Fauna Preservation Society 54, 776–83.

59. Mahmoud, S.H., Gan, T.Y., 2018. Impact of anthropogenic climate change and human activities on environment and ecosystem services in arid regions. Sci. Total Environ. 633, 1329–1344.

60. Maiti, S.K., Chowdhury, A., 2013. Effects of anthropogenic pollution on mangrove biodiversity: A review. Journal of Environmental Protection 2013, 1428–1434.

61. Malhi, Y., Gardner, T.A., Goldsmith, G.R., Silman, M.R., Zelazowski, P., 2014. Tropical forests in the anthropocene. Annual Review of Environment and Resources 39, 125–159.

62. Marshall, A.J., Lacy, R., Ancrenaz, M., Byers, O., Husson, S.J., Leighton, M., Meijaard, E., Rosen, N., Singleton, I., Stephens, S., Traylor-Holzer, K., Utami Atmoko, S.S., van Schaik, C.P., Wich, S.A., 2008. Orangutan population biology, life history, and conservation, in: Orangutans. Oxford University Press, pp. 311–326.

63. Mason, N., Ward, M., Watson, J.E.M., Venter, O., Runting, R.K., 2020. Global opportunities and challenges for transboundary conservation. Nat Ecol Evol 4, 694–701.

64. Molur, S., Walker, S., Islam, A., Miller, P., Srinivasulu, C., Nameer, P.O., Daniel, B.A., Ravikumar, L., 2005. Conservation of western hoolock gibbon (Hoolock hoolock hoolock) in India and Bangladesh. Technical Report. Zoo Outreach Organisation.

65. Morton, O., Scheffers, B.R., Haugaasen, T., Edwards, D.P., 2021. Impacts of wildlife trade on terrestrial biodiversity. Nat. Ecol. Evol. 5, 540–548.

66. Nhung, C.T.H., Le Minh, D., Anh, N.T., 2021. Modeling the distribution of the southern yellow-cheeked gibbon *Nomascus gabriellae* using maxent. Khoa Hoc Va Cong Nghe 59, 597–608.

67. Nuno, A., St. John, F.A.V., 2015. How to ask sensitive questions in conservation: A review of specialized questioning techniques. Biol. Conserv. 189, 5–15.

68. Nuttall, M.N., Griffin, O., Fewster, R.M., McGowan, P.J.K., Abernethy, K., O’Kelly, H., Nut, M., Sot, V., Bunnefeld, N., 2022. Long-term monitoring of wildlife populations for protected area management in Southeast Asia. Conserv. Sci. Pract. 4.

69. Nyhus, P.J., 2016. Human–Wildlife conflict and coexistence. Annu. Rev. Environ. Resour. 41, 143–171.

70. Pollard, E., Clements, T., Hor, N.M., Ko, S., Rawson, B., 2007. Status and conservation of globally threatened primates in the Seima Biodiversity Conservation Area, Cambodia. Wildlife Conservation Society, Cambodia Program.

71. Prakash, S., Verma, A.K., 2022. Anthropogenic activities and biodiversity threats. International Journal of Biological Innovations, IJBI 4, 94–103.

72. Rainer, H., White, A.R.T., Arcus Foundation, Lanjouw, A., 2015. Industrial Agriculture and Ape Conservation. Cambridge University Press.

73. Raum, S., 2018. A framework for integrating systematic stakeholder analysis in ecosystem services research: Stakeholder mapping for forest ecosystem services in the UK. Ecosyst. Serv. 29, 170–184.

74. Rawson, B., Hoang, M., Roos, C., Van, N., Nguyen, M., 2020. *Nomascus gabriellae*, in: IUCN Red List of Threatened Species: e.T128073282A17968950, IUCN. Accessed: 2023-8-21.

75. Rawson, B.M., Clements, T., Hor, N.M., 2009. Status and conservation of yellow-cheeked crested gibbons (Nomascus gabriellae) in the Seima biodiversity conservation area, Mondulkiri province, Cambodia, in: The Gibbons, Developments in Primatology: Progress and Prospects.

76. Rawson, B.M., Insua-Cao, P., Manh Ha, N., Ngoc Thinh, V., Minh Duc, H., Mahood, S., Geissmann, T., Roos, C., 2011. The conservation status of gibbons in Vietnam. Fauna & Flora International and Conservation International, Hanoi, Vietnam.

77. Redpath, S.M., Young, J., Evely, A., Adams, W.M., Sutherland, W.J., Whitehouse, A., Amar, A., Lambert, R.A., Linnell, J.D.C., Watt, A., Gutíerrez, R.J., 2013. Understanding and managing conservation conflicts. Trends Ecol. Evol. 28, 100–109.

78. Resasco, J., Haddad, N.M., Orrock, J.L., Shoemaker, D., Brudvig, L.A., Damschen, E.I., Tewksbury, J.J., Levey, D.J., 2014. Landscape corridors can increase invasion by an exotic species and reduce diversity of native species. Ecology 95, 2033–2039.

79. Rimbach, R., Link, A., Heistermann, M., Gómez-Posada, C., Galvis, N., Heymann, E.W., 2013. Effects of logging, hunting, and forest fragment size on physiological stress levels of two sympatric ateline primates in colombia. Conserv. Physiol. 1, cot031.

80. Sandroni, L.T., de Barros Ferraz, K.M.P.M., Marchini, S., Percequillo, A., Coates, R., Paolino, R.M., Barros, Y., Landis, M., Ribeiro, Y.G.G., Munhoes, L.P., 2022. Stakeholder mapping as a transdisciplinary exercise for jaguar conservation in the brazilian atlantic forest. Conserv. Sci. Pract. 4.

81. Scanes, C., 2018. Human activity and habitat loss: destruction, fragmentation, and degradation. Animals and human society , 451–482.

82. Sigmund, G., Ågerstrand, M., Antonelli, A., Backhaus, T., Brodin, T., Diamond, M.L., Erdelen, W.R., Evers, D.C., Hofmann, T., Hueffer, T., Lai, A., Torres, J.P.M., Mueller, L., Perrigo, A.L., Rillig, M.C., Schaeffer, A., Scheringer, M., Schirmer, K., Tlili, A., Soehl, A., Triebskorn, R., Vlahos, P., Vom Berg, C., Wang, Z., Groh, K.J., 2023. Addressing chemical pollution in biodiversity research. Glob. Chang. Biol. 29, 3240–3255.

83. Smith, J.H., King, T., Campbell, C., Cheyne, S.M., Nijman, V., 2017. Modelling population viability of three independent Javan gibbon (*Hylobates moloch*) populations on Java, Indonesia. Folia Primatol. 88, 507–522.

84. Souĺe, M.E., 1985. What is conservation biology? BioScience 35, 727–734.

85. Stark, D.J., Nijman, V., Lhota, S., Robins, J.G., Goossens, B., 2012. Modeling population viability of local proboscis monkey nasalis larvatus populations: conservation implications. Endanger. Species Res. 16, 31–43.

86. Steffen, W., Rockström, J., Richardson, K., Lenton, T.M., Folke, C., Liverman, D., Summerhayes, C.P., Barnosky, A.D., Cornell, S.E., Crucifix, M., Donges, J.F., Fetzer, I., Lade, S.J., Scheffer, M., Winkelmann, R., Schellnhuber, H.J., 2018. Trajectories of the earth system in the anthropocene. Proc. Natl. Acad. Sci. U. S. A. 115, 8252–8259.

87. Symes, W.S., Edwards, D.P., Miettinen, J., Rheindt, F.E., Carrasco, L.R., 2018. Combined impacts of deforestation and wildlife trade on tropical biodiversity are severely underestimated. Nat. Commun. 9, 4052.

88. Traeholt, C., Bunthoeun, R., Rawson, B., Samuth, M., Vutin, S., 2005. Status review of Pileated Gibbon, Hylobates pileatus, and yellow-cheeked crested gibbon, *nomascus gabriellae*, in Cambodia. Technical Report. Fauna & Flora International.

89. Tregenza, T., 1995. Building on the ideal free distribution, in: Begon, M., Fitter, A.H. (Eds.), Advances in Ecological Research. Academic Press. volume 26, pp. 253–307.

90. Treves, A., Wallace, R.B., Naughton-Treves, L., Morales, A., 2006. Co-Managing Human–Wildlife conflicts: A review. Hum. Dimensions Wildl. 11, 383–396.

91. Tunhikorn, S., Brockelman, W., Tilson, R., Nimmanheminda, U., Rantanakorn, P., Cook, R., Teare, A., Castle, K., Seal, U., 1994. Population and Habitat Viability Analysis Report for Thai Gibbons: Hylobates lar and Hylobates pileatus. Technical Report. Apple Valley, IUCN/SSC Conservation Breeding Specialist Group.

92. Vu, T.T., Tran, V.D., Giang, T.T., Nguyen, H.V., Nguyen, D.M., Nguyen, T.C., Tuyet, N.K., Doherty, P., 2016. A mark-recapture population size estimation of southern yellow-cheeked crested gibbon *Nomascus gabriellae* (Thomas, 1909) in Chu Yang Sin National Park, Vietnam. Asian Primate Journal 6, 33–42.

93. Vu, T.T., Van Dung, T., Vinh, L.Q., Nga, T.T., 2018. Using maxent to assess the impact of climate change on the distribution of southern yellow-cheeked crested gibbon (nomascus gabriellae). Management of Forest Resources and Environment 2, 131–140.

94. Wang, L., Yang, B., Bai, Y., Lu, X., Corlett, R.T., Tan, Y., Chen, X.Y., Zhu, J., Liu, Y., Quan, R.C., 2021. Conservation planning on China’s borders with Myanmar, Laos, and Vietnam. Conserv. Biol. 35, 1797–1808.

95. WCMC, U., 2021. Dataset: National protected areas of Vietnam. URL: https://data.opendevelopmentmekong.net/dataset/national-protected-areas-in-{Vxietnam.

96. Wearn, O.R., Trinh-Dinh, H., Le, Q.K., Nguyen, T.D., 2023. UAV-assisted counts of group size facilitate accurate population surveys of the critically endangered Cao Vit gibbon *Nomascus nasutus*. Oryx , 1–4.

97. Wearn, O.R., Trinh-Dinh, H., Ma, C.Y., Khac Le, Q., Nguyen, P., Van Hoang, T., Van Luong, C., Van Hua, T., Van Hoang, Q., Fan, P.F., Duc Nguyen, T., 2024. Vocal fingerprinting reveals a substantially smaller global population of the critically endangered cao vit gibbon (nomascus nasutus) than previously thought. Sci. Rep. 14, 416.

98. World Resources Institute, 2014. Global forest watch. Available at: https://www.globalforestwatch.org/. Accessed: 13 June 2023.

99. Yang, L., Chen, T., Shi, K.C., Zhang, L., Lwin, N., Fan, P.F., 2023. Effects of climate and land-cover change on the conservation status of gibbons. Conserv. Biol. 37, e14045.

