## Supplementary Tables and Figures for "Conservation of southern yellow-cheeked gibbons (*Nomascus gabriellae*) in the Anthropocene"

#### This PDF file includes:

Figs. S1 to S5
Tables S1 to S2
SI References

| Variable | Value | Explanation |
|---|---|---|
| Inbreeding in lethal equivalent | 6.29 | (1): average mammalian value; determination in primates not ethical |
| Genetic load due to recessive lethal alleles | 50 | (2): *Drosophila* estimate; determination in primates not ethical |
| EV correlation between reproduction and survival | 1 | Good years for reproduction are also good years for survival |
| EV correlation among populations | 0 | Approx. independent populations due to habitat fragmentation |
| Reproduction system | Long-term monogamy | (3) |
| First offspring age | 10 | (4): females; (3): males |
| Maximum lifespan | 30 | (3) |
| Maximum number of broods per year | 1 | (5) |
| Maximum number of progeny per brood | 1 | (3) |
| Sex ratio at birth (% males) | 52 | (4) |
| Offspring dependent for | 2 years | (4) |
| Adult females breeding | 50% of females without dependent offspring | (4): SD in offspring dependence time 50%; only females without infants breed |
| Maximum age of female reproduction | 28 | (3) |
| Maximum age of male reproduction | 30 | (3) |
| SD in % breeding due to EV | 20 | (4): offspring every $2-3$ years $(2.5 \pm 0.5$ years) |
| Distribution of broods per year | 1 brood at 100% | (5): gestation of 7 months prevents multiple broods per year |
| Number of offspring per brood per female | 1 | (5) |
| Mortality, age 0-1 | $10 \pm 3$% | (3) |
| Mortality, age 1-8 | $5 \pm 1$% | (3) |
| Mortality, age 8-10 | $15 \pm 3$% | (3): separation from group |
| Mortality, age 10+ | $5 \pm 1$% | (3) |
| Males in breeding pool | 100% | (3): all males that bred will breed |

**Table S1. Inbreeding, reproduction, and mortality of yellow-cheeked gibbons. Parameters of the population viability model. Abbreviations: EV = environmental variation, SD = standard deviation.**

| ID | Name | No Dispersal | | | Ideal Free Dispersal | | |
|---|---|---|---|---|---|---|---|
| | | Low Threats | Moderate Threats | High Threats | Low Threats | Moderate Threats | High Threats |
| 1 | Phnom Prich Wildlife Sanctuary | -0.0147 | -0.0141 | -0.0132 | -0.0345 | -0.0331 | -0.032 |
| 2 | Keo Seima Wildlife Sanctuary | -0.0142 | -0.0132 | -0.0134 | -0.033 | -0.0312 | -0.0273 |
| 3 | A Yun Pa PNR | -0.0339 | -0.0327 | -0.0319 | -0.0325 | -0.0311 | -0.0328 |
| 4 | Yok Don NP | -0.0312 | -0.0305 | -0.0241 | -0.0301 | -0.0297 | -0.0252 |
| 5 | Nui Ong NR | -0.0344 | -0.0326 | -0.0329 | -0.0294 | -0.0329 | -0.0311 |
| 6 | Chu Yang Sin NP | -0.0146 | -0.0146 | -0.013 | -0.0436 | -0.0448 | -0.0444 |
| 7 | Bi Dup-Nui Ba NP | -0.0262 | -0.0254 | -0.0211 | -0.0233 | -0.0229 | -0.0114 |
| 8 | Khanh Hoa SFE Tram Huong FC | -0.0252 | -0.0261 | -0.0192 | -0.0407 | -0.0417 | -0.0459 |
| 9 | Hon Ba NR | -0.0231 | -0.0245 | -0.0204 | -0.0355 | -0.0362 | -0.0443 |
| 10 | Phuoc Binh NP | -0.0245 | -0.0244 | -0.0228 | -0.0279 | -0.0286 | -0.0376 |
| 11 | Ninh Son SFE | -0.026 | -0.0258 | -0.0195 | -0.0225 | -0.0252 | -0.0178 |
| 12 | Nam Nung NR | -0.024 | -0.0231 | -0.0171 | -0.0227 | -0.0235 | -0.0203 |
| 13 | Ta Dung NR | -0.029 | -0.0291 | -0.0259 | -0.0291 | -0.0285 | -0.0255 |
| 14 | Quang Truc Com. | -0.0243 | -0.0254 | -0.0195 | -0.0254 | -0.0373 | -0.0586 |
| 15 | Bu Gia Map NP | -0.0155 | -0.0154 | -0.0148 | -0.0411 | -0.0548 | -0.0897 |
| 16 | Cat Tien NP | -0.0148 | -0.0142 | -0.0124 | -0.0442 | -0.0459 | -0.0458 |
| 17 | Loc Bac SFE | -0.0325 | -0.0325 | -0.0312 | -0.0082 | -0.0094 | -0.003 |
| 18 | Dong Nai NR | -0.0284 | -0.0284 | -0.0231 | -0.0118 | -0.0125 | 0.0045 |
| | **Metapopulation** | -0.0149 | -0.015 | -0.0141 | -0.034 | -0.0351 | -0.0323 |

**Table S2. Stochastic mean growth rate. Results of the population viability modelling for populations in Table 1 with known population sizes. The population viability is performed for various levels of deforestation and hunting (threat levels given in Table 2) and for variable dispersal between populations (no dispersal or ideal free dispersal). The stochastic mean growth rate is determined for each scenario and each population by running the stochastic simulation 100 times and measuring the growth rate in each simulation (see Methods for details). A negative mean stochastic growth rate indicates a declining population.**

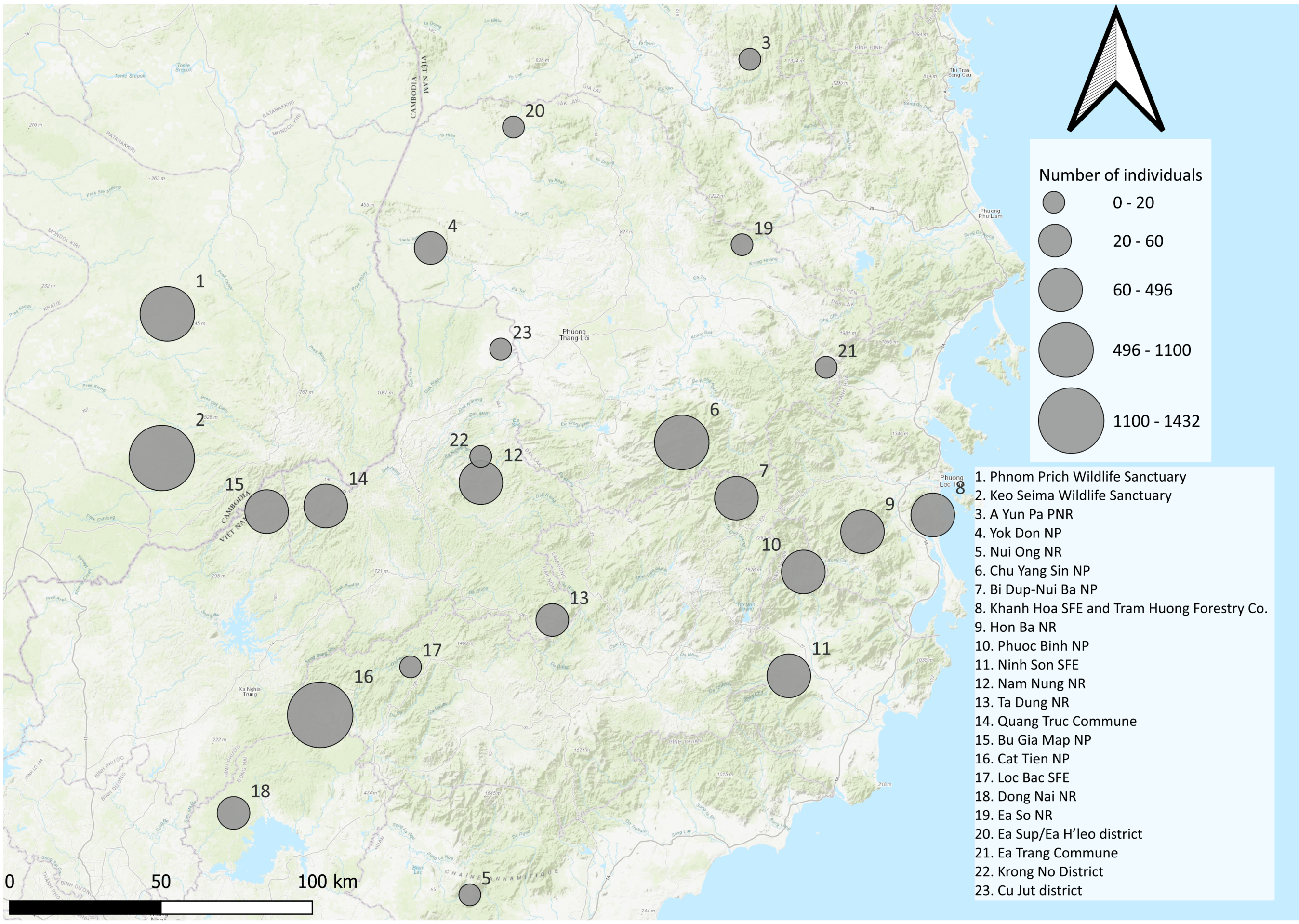


**Fig. S1. Population sizes of yellow-cheeked gibbons in Vietnam and Cambodia.** The population sizes of areas in Fig. 1 and Fig. 2a are plotted on a map as grey circles with a size that reflects the population size.

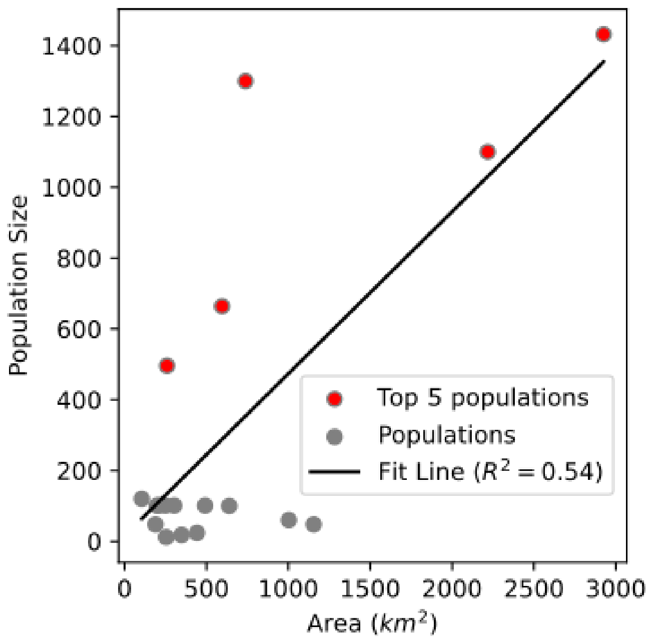


**Fig. S2. Correlation between population size and area.** The sites in Fig. 2a with known area (x-axis) and population size (y-axis) are represented by grey dots. The five sites with the largest reported population (dots with red centre) are (from largest to smallest): Keo Seima Wildlife Sanctuary, Cat Tien National Park, Phnom Prich Wildlife Sanctuary, Chu Yang Sin National Park and Bu Gia Map Nation Park. A line of best fit is plotted ($R^2$=0.54), $\text{Population Size} = 0.46 \times \text{Area}\ [\text{km}^2] + 15.82$. All five largest populations lie above the line of the best fit.

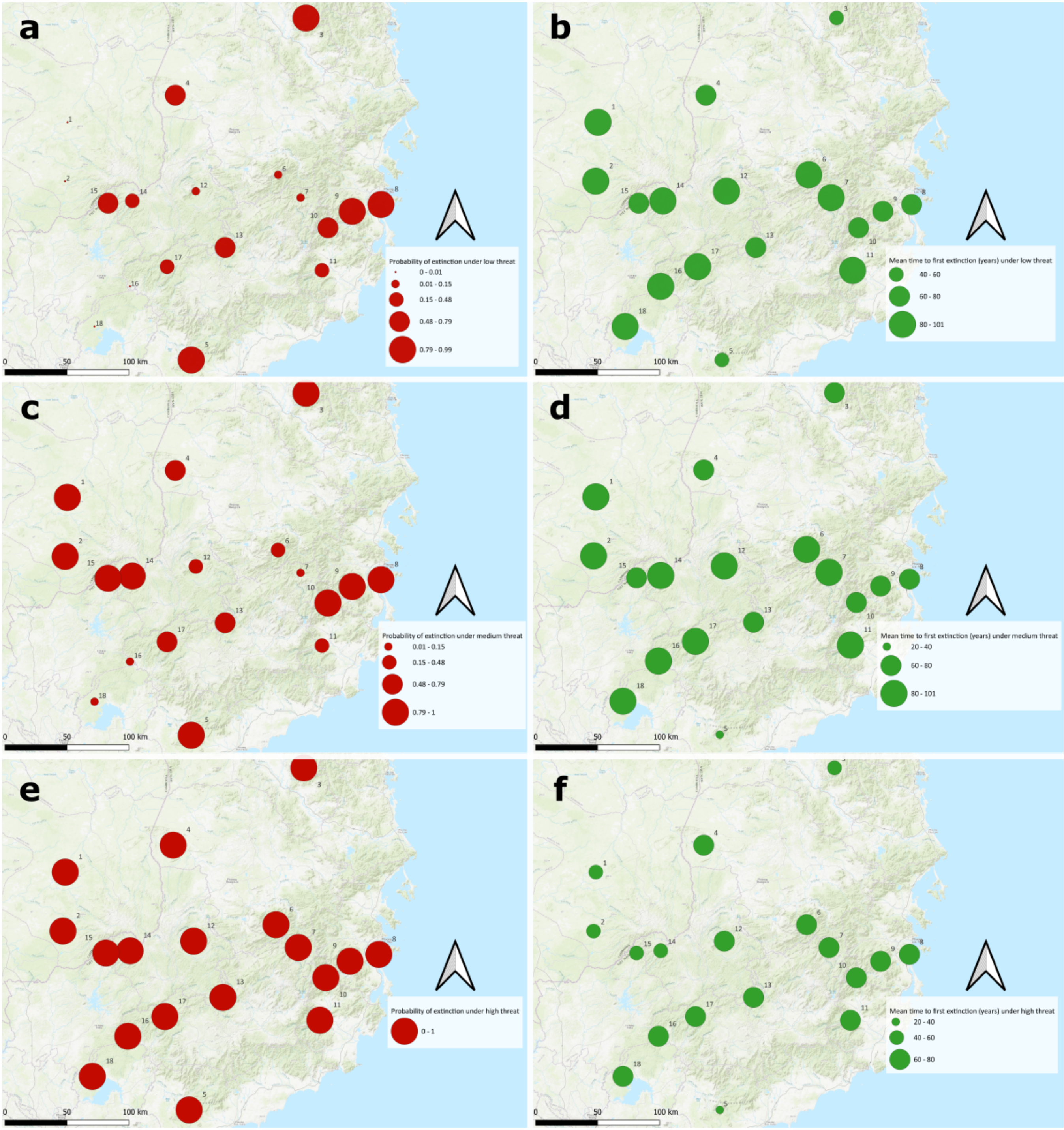


**Fig. S3. Probability of extinction and mean time to first extinction for yellow-cheeked gibbon populations under variable levels of deforestation and hunting.** Data in Fig. 2d and Fig. 2f visualized on a map. Probability of extinction is denoted by red circles of proportionate sizes in panels (a, low threat, c, moderate threats, e, high threats). Mean time to first extinction is denoted by green circles of proportionate sizes in panels (b, low threat, d, moderate threats, e, high threats). The levels of threats (namely deforestation and hunting) are given in Table 2. The model with the ideal free dispersal is considered.

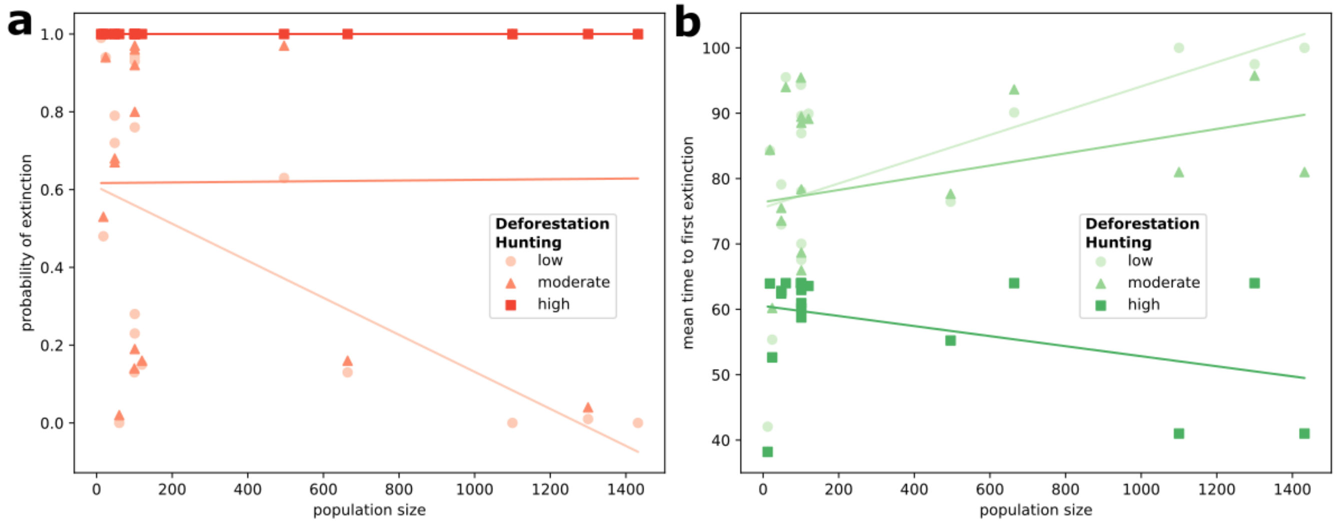


**Fig. S4. Correlation between population size and extinction for various threat levels. (a)** Correlation with extinction probability. Points correspond to sites, the line of the best fit is obtained by linear regression. The model with dispersal and different levels of deforestation and hunting in Table 2 is considered: low threat (light red, circle, fitted $R^2$=1), medium threat (red, triangle, fitted, $R^2$=0.0001), high threat (dark red, square, fitted $R^2$=0.3427). **(b)** Correlation with the mean time to the first extinction. Points correspond to sites, the line of the best fit is obtained by linear regression. The model with dispersal and different levels of deforestation and hunting in Table 2 is considered: low threat (light green, circle, fitted $R^2$=0.3068), medium threat (green, triangle, fitted, $R^2$=0.0896), high threat (dark green, square, fitted $R^2$=0.1673). In both panels, the largest populations exhibit the largest improvement as deforestation and hunting are reduced.

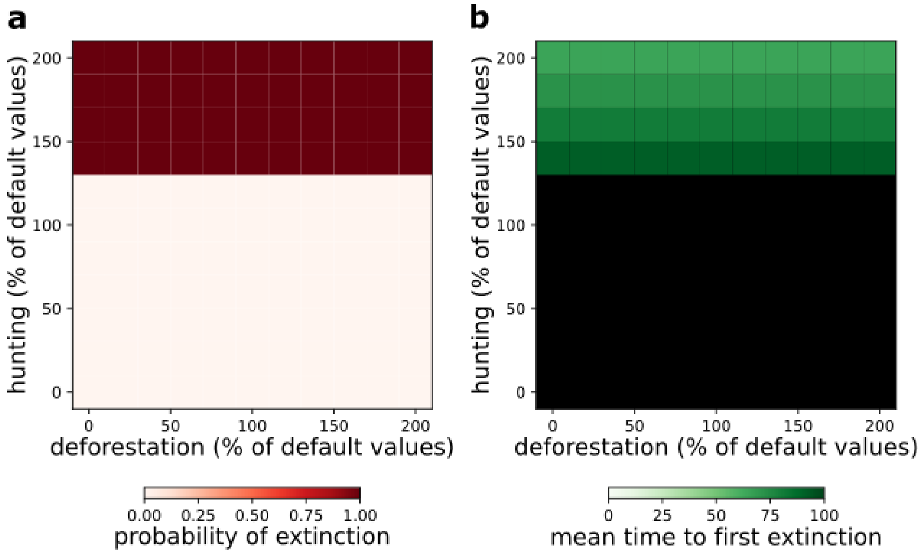


**Fig. S5. Sensitivity of metapopulation to hunting and deforestation.** Probability of extinction (red heatmap, a) and mean time to first extinction (measured in years, green heatmaps, b) are determined for variable levels of deforestation rate (x-axis) and hunting rate (y-axis). The default values of deforestation and hunting rates that correspond to 100% along these axes are given by the moderate values in Table 2. Correspondingly, the low (resp. high) levels of deforestation and hunting in Table 2 correspond to a point with 50% (resp. 200 %) deforestation and hunting rates. The overall pattern, exhibiting a strong sensitivity to hunting, is similar to the pattern of large sites in Fig. 3.

### References


1. JJ O'Grady, et al., Realistic levels of inbreeding depression strongly affect extinction risk in wild populations. *Biol. Conserv.* **133**, 42–51 (2006).
2. MJ Simmons, JF Crow, Mutations affecting fitness in *Drosophila* populations. *Annu. Rev. Genet.* **11**, 49–78 (1977).
3. C Traeholt, R Bunthoeun, B Rawson, M Samuth, S Vutin, Status review of pileated gibbon, hylobates pileatus, and yellow-cheeked crested gibbon, *nomascus gabriellae*, in Cambodia, (Fauna & Flora International), Technical report (2005).
4. P Fan, et al., Reproductive parameters of captive female northern white-cheeked (*nomascus leucogenys*) and yellow-cheeked (*nomascus gabriellae*) gibbons. *Int. J. Primatol.* **42**, 49–63 (2021).
5. NTT Hien, NT Binh, An overview of the yellow-checked gibbon (nomascus gabriellae thomas, 1909). *J. Thu Dau Mot Univ.* pp. 207–213 (2024).
